# Phosphorylation-dependent binding of the Adenomatous Polyposis Coli protein to β-TrCP1 regulates β-catenin destruction

**DOI:** 10.64898/2026.09.16.751897

**Authors:** Darius M. Johnston, Crissy L. Tarver, Ryan Leib, Mardo Kõivomägi, Fang Liu, Christopher Adams, William I. Weis

**Affiliations:** Department of Molecular Cellular Physiology, Stanford University School of Medicine, Stanford, CA; Department of Structural Biology, Stanford University School of Medicine, Stanford, CA; Vincent Coates Foundation Mass Spectrometry Laboratory, Stanford University, Stanford, CA; Laboratory of Biochemistry and Molecular Biology, National Cancer Institute, NIH, Bethesda, MD 20892, USA

## Abstract

Wnt signaling controls cellular development by regulating levels of β-catenin, a dual transcription factor and adhesion protein. The tumor suppressor gene Adenomatous Polyposis coli (APC) is a negative regulator of Wnt signaling that acts by interacting with multiple proteins in the destruction complex to drive β-catenin degradation. Mutations in APC cause deregulation of Wnt/β-catenin signaling contributing to the development of several types of cancer, especially colorectal cancers. Tumors are not null mutant for APC—instead they accumulate truncated proteins. Many APC proteins are truncated between the second and third 20-amino acid repeats, a region known as the β-catenin inhibition domain (CID), which is essential for β-catenin destruction, but its mechanism of action is unclear. Here we find that cyclin dependent kinases and GSK3β phosphorylate the CID region of APC *in vitro*. This allows the APC CID to bind to the E3 ubiquitin ligase component β-TrCP1 in a phosphorylation dependent manner and this in turn regulates the rate of β-catenin ubiquitylation. β-TrCP1 binding to the phosphorylated CID favors the retention and stabilization of the SCF^β-TrCP1^ E3 ligase in the destruction complex and ensures that SCF^β-TrCP1^ remains available for β-catenin ubiquitylation. These findings provide a mechanism for the vital role of APC’s CID region in β-catenin destruction.

## Introduction

The Wnt/β-catenin signaling pathway performs critical roles in development and tissue homeostasis^1–4^. Binding of a secreted Wnt growth factor to its cell surface receptors results in the stabilization of the transcriptional co-activator β-catenin and activation of target genes that control cell growth and differentiation^5^. In the absence of a Wnt signal, β-catenin is phosphorylated, ubiquitylated, and targeted for proteasomal destruction^5^. These negative regulatory activities occur in a “destruction complex” that contains the proteins Axin and Adenomatous Polyposis Coli (APC), the kinases GSK3 and CK1, and the SCF^β-TrCP^ E3 ubiquitin ligase^6^ (Figure 1A). Axin binds to APC, β-catenin, GSK3 and CK1, thereby acting as a scaffold for phosphorylation of β-catenin that creates the phosphoserine recognized by β-TrCP1. Mutations in destruction complex components are associated with several cancers, notably colorectal cancer, due to inappropriate stabilization of β-catenin^7–9^.

**Figure 1:**
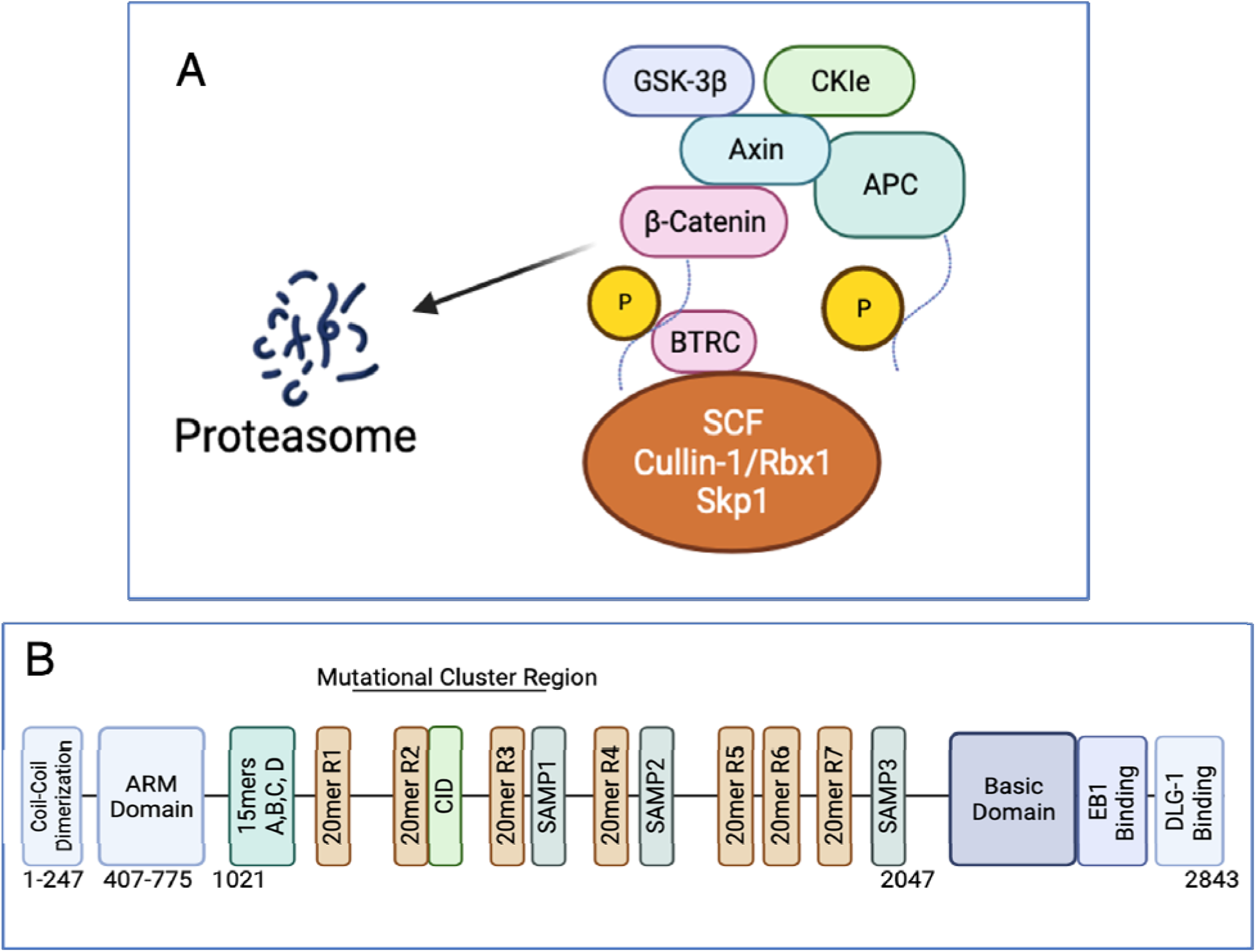
A) A cartoon overview of the Destruction Complex showing the core components including two kinases, Ck1 and GSK3, two scaffolds, Axin and APC, and the E3 ligase SCF(BTRC). Axin scaffolds the phosphorylation of APC, which increases the affinity for Beta-catenin, allowing Axin to phosphorylate Beta-catenin. This creates a phospho-degron that is recognized by SCF(BTRC), which ubiquitinates Beta-catenin such that it is destroyed by the proteasome. B) An overview of the domain structure of APC. This includes an N terminal coil-coil domain that induces dimerization, an armadillo domain, four beta-catenin binding 15mer motifs, six beta-catenin binding 20mer repeats, one 20mer that does not bind beta-catenin, three Axin binding SAMP repeats, a basic domain important in microtubule regulation, an EB1 binding domain for localization of the protein and a DLG1 binding domain thought to retain APC during cell polarity functions. The N terminal armadillo domain has shown to be absolutely essential for regulating actin during cell migration and more recently to be necessary for correct destruction complex activity. The mutational cluster region of colorectal cancer patients is annotated above.

APC plays roles in cell-cell adhesion^10^, cytoskeletal regulation^11–13^, and cell cycle regulation^14,15^, but is best known as a negative regulator of Wnt/β-catenin signaling, as it is a critical component of the destruction complex^16–18^. Human APC is a 2843 amino acid protein that has a N-terminal dimerization domain, an armadillo repeat domain, a central region that contains four homologous 15-and seven 20-amino acid repeats (15R A-D and 20R 1-7) that bind β-catenin, and three SAMP repeats that bind to Axin19. It also contains a C-terminal region that includes EB1 binding, microtubule regulating and actin-binding activities as well as a C-terminal peptide that binds DLG1^20–22^ (Figure 1B).

APC’s disordered nature, large size and multiple β-catenin binding sites in the central region have complicated mechanistic studies. The multiple β-catenin binding repeats are strengthened by phosphorylation of APC and are thought to sequester β-catenin, thereby preventing inappropriate activation of Wnt target genes prior to β-catenin destruction^23,24^. Notably, while the 15mers appear dispensable for β-catenin destruction, at least one β-catenin binding 20mer is required^25^. In addition, the different affinities displayed, as well as their change in affinity is based on the phosphorylated states of the 20mers, provide a range of affinities to β-catenin that may help to fine tune the response for β-catenin destruction (apart from 20mer R2, which does not bind to at all)^26^. At a higher level of organization, there is evidence that multiple copies of APC, by virtue of its ability to dimerize and bind multiple Axin molecules, may contribute to higher order phase behavior, including phase separated material^27^. Certain properties of APC, like its multivalent character and intrinsically disordered regions, indicates APC may form a condensate^28^. One hypothesis is that high local concentrations are necessary for efficient phosphorylation and ubiquitylation activities in the destruction complex^29^.

One unusual feature of APC as a tumor suppressor is that tumors are not null mutant for APC. Instead, all express truncated APC proteins due to the crucial role of the N terminal region in cell migration and division^11^. Despite the complex structure of APC, most oncogenic truncations occur in a minimal and highly conserved region called the mutational cluster region (MCR), which includes 20mer repeats R1 through R3 as well as the SAMP1 region^30,31^ (Figure 1B). These truncations are responsible for disrupting β-catenin destruction. The MCR contains at least one β-catenin-binding 20mer repeat and the highly conserved, non-binding 20R2 which is required for β-catenin ubiquitylation^32^. It also includes the sequence between 20R2 and R3, which is strongly conserved and essential for β-catenin destruction but does not bind β-catenin^33^. The functional role of this region, designated the Catenin Inhibitory Domain (CID), also called Region B in *Drosophila melanogaster*, has eluded mechanistic definition. In one model, the CID regulates a putative interaction between Axin and the APC ARM repeats such that phosphorylation of the CID disrupts this interaction, which then allows Axin to scaffold β-catenin phosphorylation^34^. It has also been proposed that the CID inhibits the DUB USP7, which would otherwise de-ubiquitylate β-catenin^35^, although more recent findings have challenged this model^36^. The suggestion that the CID inhibits the activity of a phosphatase needed to remove β-catenin from its tight-binding phosphorylated repeats^37^ appears incompatible with the insensitivity of phosphorylated R3 to phosphatase over many hours^24^. Furthermore, the absence of a phosphatase in a reconstituted system did not impair ubiquitylation or proteasomal destruction of β-catenin^38^.

While most colorectal tumors carry bi-allelic mutations in APC, other tumors carry β-catenin mutations at residues 33/37/41/45. These prevent proper phosphorylation, which is critical for recognition by F-BOX protein β-TrCP1^39^. This region binds to the E3 SCF^β-TrCP^ ubiquitin ligase and significantly accelerates ubiquitin transfer to β-catenin, consistent with the association of β-TrCP with the destruction complex^40,41^.

Given its key role in APC function, it is critical to define the functions of the R2/CID region. Here we demonstrate that the R2-CID region of APC binds to β-TrCP1 in a phosphorylation-dependent manner, and this significantly increases the rate of ubiquitin transfer to β-catenin. These effects are dependent upon phosphorylation of the R2-CID by GSK3β and, unexpectedly, cyclin-dependent kinases (Cdks). These data indicate that phosphorylation dependent binding enables retention and stabilization of the critical E3 ligase to the destruction complex. Phosphorylation of R2-CID by Cyclin/Cdks indicates a fundamental link of β-catenin destruction to the cell cycle.

## RESULTS

### β-TrCP1 and Cdks associate with the CID region of human APC

We employed a rapid capture mass spectrometry screen to search for specific binding partners to APC’s 20mer R2-CID region (Figure S1A). HEK 293 cells were grown in media lacking leucine and methionine for 16 hrs, followed by growth in media containing photo-activatable diazirine derivatives of leucine and methionine for 24 hrs^42^. The cells were then transfected with a streptavidin-binding protein (SBP)-and HA-tagged APC construct spanning the R2-CID region, or a longer construct that contained the R2-CID-R3 sequence (Figure 1B). The latter construct allowed us to distinguish specific R2-CID partners from those that might bind through β-catenin, which binds strongly to the 20mer R3 but not to R2^24,26^. After 24 hrs, the cells were treated with the proteasome inhibitor MG132 for 2 hrs to slow flux through the destruction complex by preventing degradation of ubiquitinated β-catenin. The cells were washed with PBS and exposed to ultraviolet light to activate crosslinking (Figure S1B). The APC constructs were captured on streptavidin resin, fractionated, and cross-linked bands were submitted for LC-MS/MS analysis.

We first validated the APC R2-CID-R3 sample to ensure that known destruction complex partners were present. Axin and β-catenin were found in this sample but not in the R2-CID, indicating these proteins are specific partners of the R3 region. Other non-crosslinked proteins of interest are noted in Table S1. These data served as a useful check on the recovery of proteins in the R2-CID region, which was the principal focus of this study.

Several approaches were used to detect peptides crosslinked to only the R2-CID but not R2-CID-R3. Byonic X-link (Proteometrics) allows for a ‘wild card’ search that allows for a wide variation of molecular weights added to specified residues. We analyzed given these variable modifications to each side chain of both the donating leucine or methionine, and the accepting residues to determine if a cross-link occurred (See Methods). This was done serially and with dummy data sets to validate the presence of cross-linked peptides. When we separated the specifically cross-linked peptides from the rest of those found in the screen, it revealed several specific hits (Table S2). These data revealed interactions between APC and β-TrCP1/Skp1, but not with β-catenin (Table S2). More surprisingly, both non-crosslinked and crosslinked data included several cyclins and cyclin-dependent kinases (Cdks) interacting with the R2-CID (Table S1 and S2), which prompted our investigation of this interaction.

Given that these analyses used the human APC sequence, we asked whether these findings are generalized given the conservation of the R2-CID region. While the entire R2-CID region is highly conserved in bilaterian animals, the sequence corresponding to the human motif ^1410^CSGMVS^1415^, is also strongly conserved from Hydra to chordates. Intriguingly, β-TrCP recognizes and binds to substrates by means of a conserved motif, DpSGXXpS, where pS is phosphorylated serine (Figure S3A). The phosphoserines at positions *n* and *n+4*, as well as a conserved glycine at *n+1*, are essential for the binding of β-catenin and many other substrates^43^ (Figure 2A).

**Figure 2:**
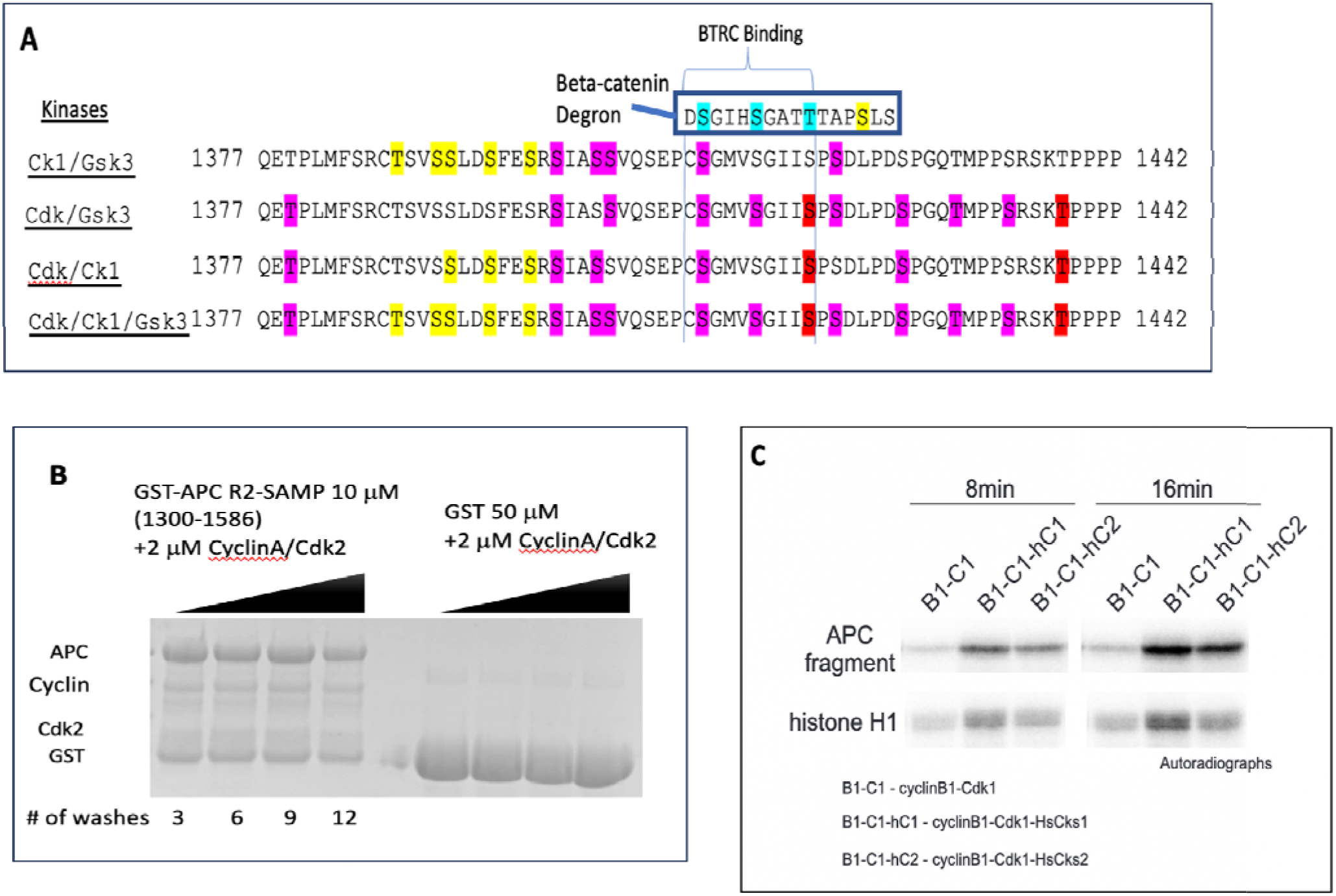
A) Phosphorylation patterns of the APC CID Region A sequence alignment showing the different phosphorylation patterns in the CID region given by different kinase combinations. Magenta denotes sites identified in this study. Red denotes Cdk sites and yellow shows previously identified sites. (Rubenfeld, Polakis) (See also figure S2A). The N terminal region of Beta-catenin is shown above with the BTRC degron sites shown in blue. B) APC can bind to CyclinA/Cdk2. A GST pulldown showing that recombinant APC 1300-1586 is able to bind recombinant CyclinA/Cdk2 with free GST as a control. 20uL of Glutathione beads were washed repeatedly in alternating high (1M NaCl) and low (100mM NaCl) salt buffers in 1mL washes with 5min incubations. C) APC residues 1300-1586 phosphorylated with CyclinB/Cdk1 and Cks1 or Cks2 + GSK-3β and histone H1 as a positive control. Cks increases phosphorylation of the APC CID region; p32 labeled ATP was incubated with indicated kinase/protein combination to determine if Cks1 or Cks2 altered Cdk phosphorylation of APC.

Thus, many sequences in this part of the CID, including those in the best studied animals, mammals and Drosophila, fit the core β-TrCP1 binding motif at the *n*, *n+1* and *n+4* positions. A second, overlapping match to this motif is also found in the CID region of human APC, which allows Cdk phosphorylation at 1419 to prime phosphorylation of both 1415 and 1411 by GSK3. ^1410^CSGMVSGIISP^14^^20^ (Figure 2A). Although the β-TrCP consensus shows an aspartate at position *n-1*, Alphafold3 models based upon the crystal structures of β-catenin bound to β-TrCP^44,45^ suggested that a cysteine and possibly a valine can be accommodated in this position (Figure S1C). Furthermore, β-TrCP can recognize the degron site TpSGXXpS of cyclin F once it is phosphorylated by CK2^46^. These observations led us to hypothesize that the APC CID regions 1410-1415 and/or 1414-1419 could bind to β-TrCP when the critical serine residues are phosphorylated.

The phosphorylation motif for GSK3 recognizes includes a Ser or Thr four residues C-terminal a Ser or Thr that was previously phosphorylated by another kinase, a process known as priming^48^. For β-catenin, CK1 provides the initial priming by phosphorylating S45, and successive GSK3 phosphorylation at sites S41, S37, and S33 to generate the phosphoserines^47,48^. In the APC CID region, GSK3 phosphorylation of residues S1411 and S1415 would require priming of S1419, which is not a GSK3 or CK1 site. However, S1419 is followed by P1420 (^1410^CSGMVSGIISP^1420^). The sequence around S1419 is a potential phosphorylation site for Cdks, which phosphorylate a Ser or Thr that is followed by Pro^49^. This led us to the hypothesis that Cdk’s could be interacting with APC CID region to prime GSK3.

Since Cdks were found in the mass spectrometry screen, we tested the hypothesis that the R2-CID region of APC interacts with Cyclin A/Cdk2. A GST pulldown showed an interaction between recombinant APC (1300-1585) with recombinant Cyclin A/Cdk2 with free GST as a control (Figure 2B). We next hypothesized that the R2-CID could be sequentially phosphorylated by a combination of CyclinA/Cdk2 or Cyclin B/Cdk1 and their regulatory subunits Cks1 or Cks2. Cyclins A and B produce nearly the same active confirmation of Cdk1 and Cdk2, but Cdk1/CyclinB was tested due to having higher activity for enzymology experiments. To test whether CDKs can phosphorylate the CID, we used the same APC construct (1300-1585) for *in vitro* phosphorylation by Cyclin B/Cdk1 and their regulatory subunits Cks1 or Cks2, with histone H1 used as a positive control. Autoradiographs revealed robust R2-CID phosphorylation, with Cks1 and Cks2 increasing phosphorylation of this APC fragment (Figure 2C). This suggests that this region of APC is a bona fide substrate of Cyclin/Cdk/Cks complexes and can interact with this APC construct.

CyclinA and CyclinE bind substrates with an (R/K)-X-L motif, while CyclinB binds more specifically to (R/K)-X-L-Φ or (R/K)-X-L-X-Φ where X is any amino acid and Φ is a large hydrophobic residue^50^. The R2/CID region of APC contains a related motif, ^1345^A**RHK**AVE**F**^1354^, which most closely resembles the cyclin binding site on P53, ^377^S**RHK**KLM**F**^385^, with the sole change of a Leu to Val in the non-X positions. We used Alphafold3 to assess whether this APC motif might bind CyclinA in the same manner when compared to the CyclinA/Cdk2/P53 structure (PDB:1H26). This produced a high confidence model that suggests APC could interact with CyclinA in a manner similar to P53 (Figure S1C), where there was the loss of only one interaction due to the Leu=>Val substitution. This would provide a potential mechanism by which Cyclin binding to APC could recruit CDKs to phosphorylate the CID to prime GSK3.

### APC R2/CID binds to β-TrCP1 in a CDK/GSK3β-phosphorylation-dependent manner

Ranes et al. 2021, found that APC physically interacts with the SCF^β-TrCP1^ E3 ligase complex^39^. We examined whether this interaction would still occur if we limited APC to the CID region, and whether it depended on phosphorylation. To do so, we asked whether they would co-elute during analytical gel filtration experiments. To prepare samples, APC R2-SAMP1 (1342-1585) was affinity purified and treated with lambda phosphatase to remove any non-specific phosphates. This protein was then re-purified to obtain a pure sample lacking any residual phosphorylation or phosphatase. A sample of this protein then underwent *in vitro* phosphorylation using Cyclin A/Cdk2/Cks2 and GSK3β. After *in vitro* phosphorylation, this protein was further purified over MonoQ resin to separate phosphorylated from non-phosphorylated subpopulations to obtain homogenous samples and remove the kinases. Phosphorylation or non-phosphorylation of these samples was verified using a Pro-Q Diamond Phosphoprotein gel stain (ThermoFisher) gel after cleavage of GST-tags (Figure S4a). This stain reacts strongly to phosphorylated residues to indicate the presence or absence of phosphorylation on proteins in SDS-PAGE.

We next used these samples in analytical gel filtration experiments along with β-TrCP1/Skp1. The elution peak for non-phosphorylated APC centered around 8 mL, while the elution peak for APC phosphorylated with Cyclin A/Cdk2/Cks2 + GSK3βG centered around 9 mL. The elution peak for β-TrCP1/Skp1 alone was centered at 18 mL. When we mixed APC phosphorylated with Cyclin A/Cdk2/Cks2 + GSK3β and β-TrCP1/Skp1, we saw a single elution peak centering around 13 mL, suggesting that they interact quantitatively (Figure 3A). To confirm this, fractions from this one peak were analyzed by SDS-PAGE, and both APC and β-TrCP1/Skp1 were present (Figure 3B). We confirmed that elution of β-TrCP1/Skp1 was not altered by non-phosphorylated APC as a mixture of these proteins resulted in two elution peaks. SDS-PAGE revealed that each peak contained only one of the two proteins (Figure 3C). These data suggest that phosphorylation of R2/CID by Cdk/GSK3β allows binding to β-TrCP1/Skp1. A similar result was not seen in gel filtration experiments mixing APC phosphorylated with CK1 + GSK3β and β-TrCP1/Skp1(dns).

**Figure 3:**
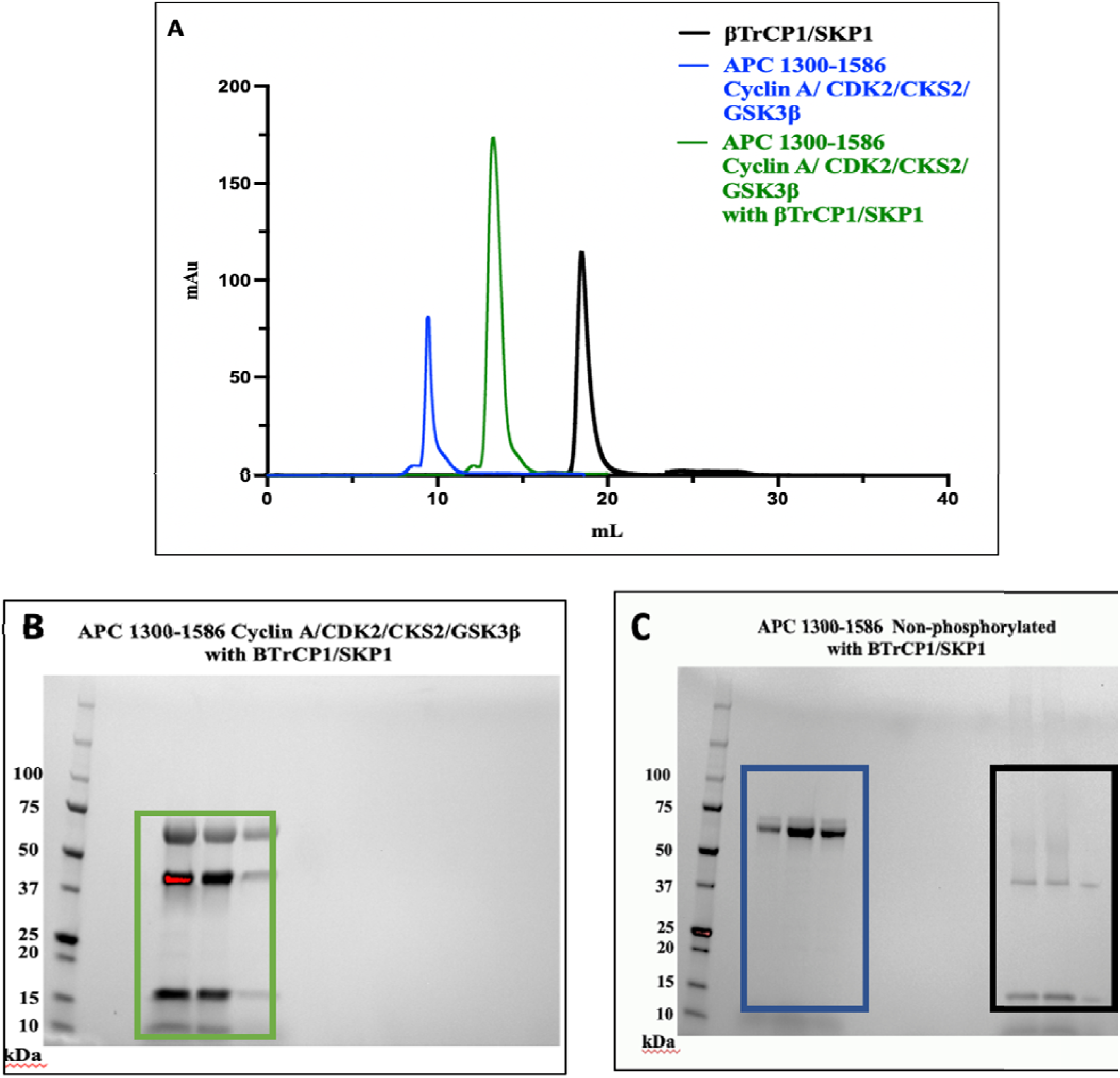
Size exclusion chromatography shows co-elution of BTRC/SKP1 with APC residues 1300-1586 phosphorylated with CyclinA/Cdk2/Cks2 + GSK3β. A) Chromatograph showing elusion peaks for BTRC/SKP1 alone (black), APC residues 1300-1586 phosphorylated with CyclinA/Cdk2/Cks2 + GSK-3β alone (blue), and a sample containing a combination of these two proteins (green). B) Fraction containing protein from the green peak analyzed on SDS-PAGE. C) Sample of non-phosphorylated APC residues 1300-1586 mixed with BTRC/SKP1 and resolved by size exclusion chromatography. Fractions from each peak analyzed on SDS-PAGE.

To measure the quantitative affinity and kinetic aspects of this interaction, we used Biolayer interferometry (BLI). We first tried used biotinylated Avi-β-TrCP1/Skp1 and APC CID residues 1300-1586 (the region used for the co-elution experiments). However, this segment of APC proved too unstable for these measurements, so a smaller region of the APC CID (1342 – 1438) was recombinantly expressed and purified as above to use in these studies. No binding was detected with lambda phosphatase-treated (non-phosphorylated) APC. In contrast, the Cyclin A/Cdk2/Cks2 + GSK3β phosphorylated material bound with a K_D_ of 7.65 µM (Figure 4A & B), consistent with other known β-TrCP degradation targets^51^. As a positive control, we produced CK1 + GSK3β phosphorylated β-catenin as a substrate to determine binding to biotinylated Avi-β-TrCP1/Skp1. The phosphorylated β-catenin bound to β-TrCP1/Skp1 with a K_D_ of 0.5 µM, a similar affinity as previously reported^45^ (Figure 4C). Intriguingly, the 15x fold weaker affinity of the phosphorylated APC CID to Avi-β-TrCP1/Skp1relative to β-catenin binding to Avi-β-TrCP1/Skp1arises from both a 205x smaller on-rate constant and a 13x smaller off-rate (Figure 4C). Thus, phosphorylation of the R2/CID region leads to binding to β-TrCP1/Skp.

**Figure 4.**
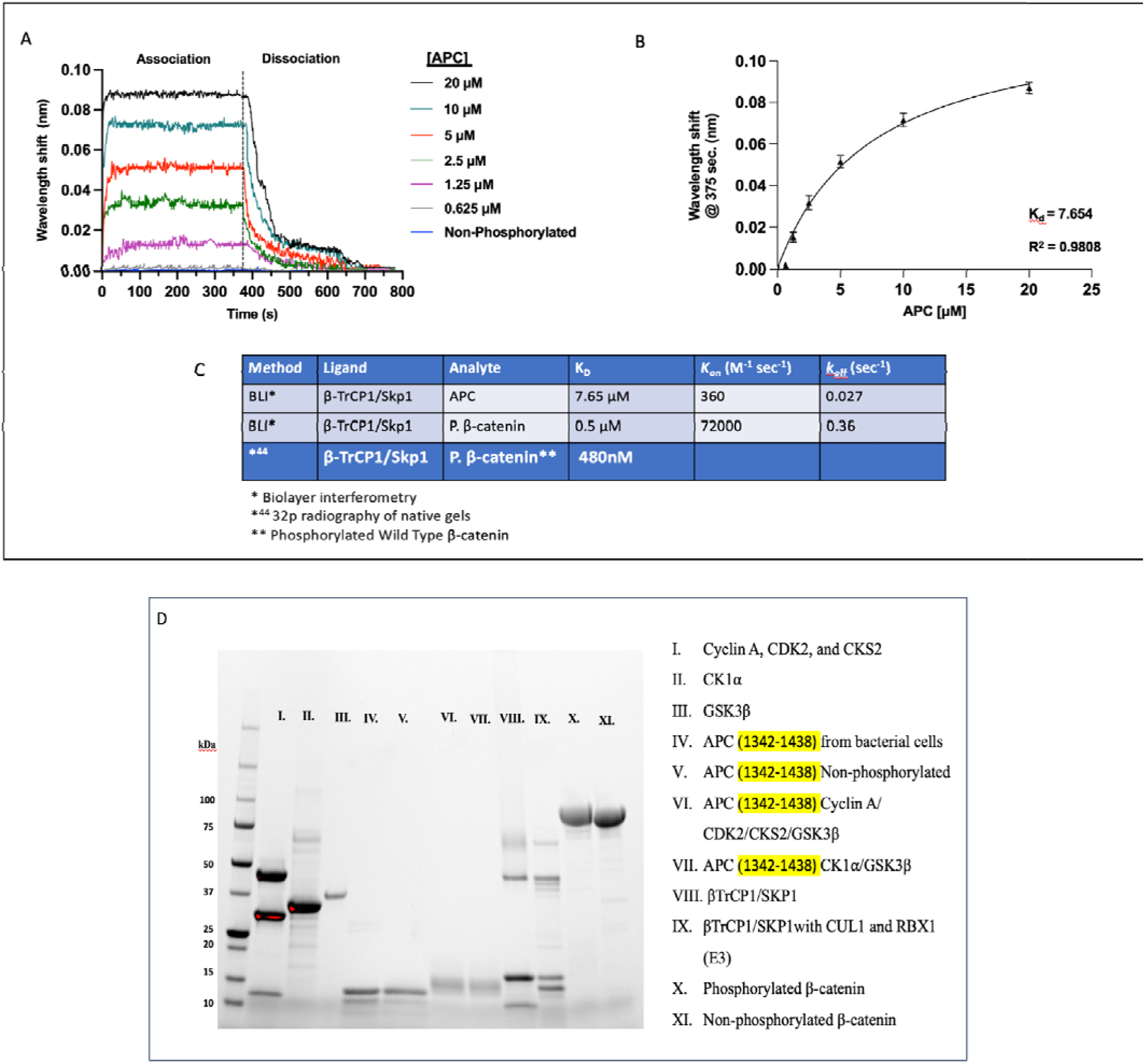
BTRC/SKP1 interacts with APC CID in a phosphorylation-dependent manner. A) Binding of BTRC/SKP1 to the CID region of APC was monitored using biolayer interferometry (BLI). Association and dissociation curves of biotinylated BTRC/SKP1immobilized on streptavidin biosensors are shown binding to varying concentrations of APC. Samples of Cdk2/GSK3 phosphorylated APC and non-phosphorylated APC were examined. B) Wavelength shifts at 375 s (plateau) of each APC CID concentration was graphed with data points shown as mean ± SEM from n=3 experiments. Data fitted to the non-linear equation Y=B_max_*X/(K_d_+X), where B_max_ is the maximum shift in wavelength, and the K_d_ and R^2^ values were determined. C) Table shows kinetic data for BTRC/SKP1 to phosphorylated APC and phosphorylated β-catenin. β-catenin was purified and underwent *in vitro* phosphorylation with Ck1/GSK-3β. The binding of BTRC/SKP1 to phosphorylated β-catenin was examined by BLI and compared to previously published results obtained by p32 radiographic analysis of native gels^44^.

### Phosphorylated APC R2-R3 enhances ubiquitylation of β-catenin

To assess the effect of phosphorylated APC binding to β-TrCP1/Skp1 on ubiquitin transfer to β-catenin, we created an APC construct spanning R2-R3 (residues 1340-1532) (Figure 1C). We compared samples of non-phosphorylated APC with samples that were phosphorylated *in vitro* by purified CyclinA/Cdk2/Cks2 + GSK-3 (“CCC”; Figure 5A). Purified β-catenin that had been CK1 and GSK-3 phosphorylated was fluorescently labeled with carboxy-rhodamine^52^, and ubiquitin protein was trace fluorescently labeled with Texas Red. Each reaction was performed using the minimal ubiquitin transfer machinery, including UBA1(E1), Ubc5(E2), CUL1, Rbx1(E3), Skp1, β-TrCP1, APC, β-catenin, and Ubiquitin. The pre-purified phosphorylated APC R2-R3 and β-catenin eliminated the need for the addition of free kinases in the reactions such that only one enzyme was consuming ATP in the reaction.

**Figure 5).**
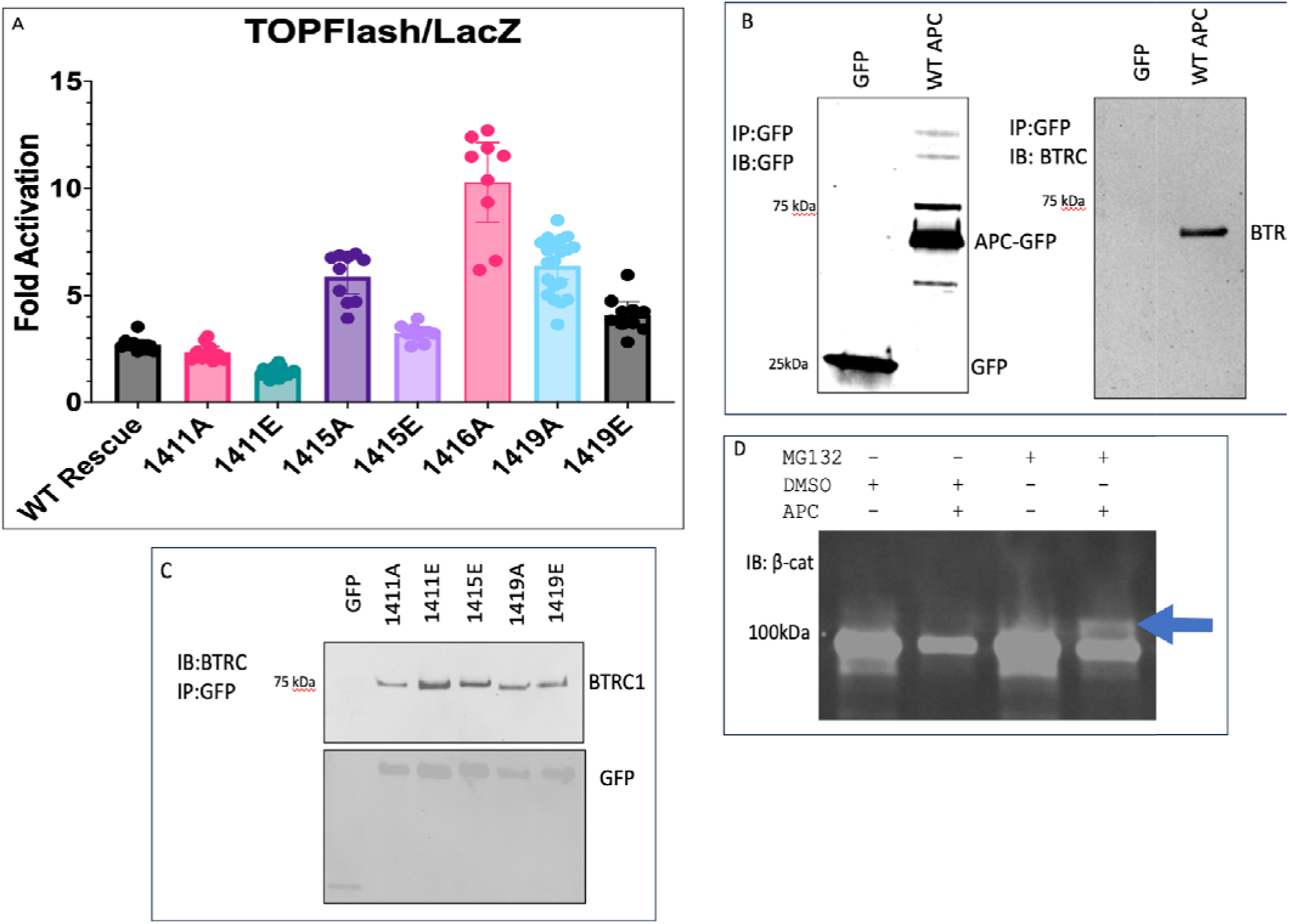
A) Point Mutations of the BTRC Binding Motif in APC affect Wnt Signaling TOPFLASH assay showing the effect of disrupting the BTRC/APC interface residues, either by mutating priming serine 1419, GSK3 sites 1411 and 1415 or the conserved glycine 1416 in the BTRC binding motif. This was compared to a WT rescue. b) APC can immuno-preciptiate BTRC in 293 cells. CRISPR truncated APC 293 cells were transfected with GFP-APC rescue mutants and immunoprecipitated by an anti-GFP nanobody resin. Then were then serially stained as indicated. c) Mutants disrupting kinase sites were tested for altering. D) MG132 Treatment shows distinct ubiquitin band for Beta-catenin in the presence of APC-GFP. APC Min cells transfected with either GFP or GFP-APC WT were treated with MG132 overnight. Lysate was then blotted against Beta-catenin

Phosphorylated APC R2-R3 significantly enhanced the rate of ubiquitin transfer to β-catenin in comparison to non-phosphorylated APC R2-R3 (Figure. 6). Previous work revealed that phosphorylation of APC R3 by CK1 + GSK-3 substantially elevates its binding affinity to β-catenin^24,26^. However, the non-CK1 treated construct still binds to β-catenin with a K_D_ of 200 nM. Since the APC R2-R3 in this assay was not phosphorylated with CK1, CK1 phosphorylation of APC is not essential for enhancing β-catenin ubiquitination. Thus, these data demonstrates that phosphorylation of R2-R3 by CDKs and GSK3 increase the rate of ubiquitin transfer to β-catenin.

**Figure 6.**
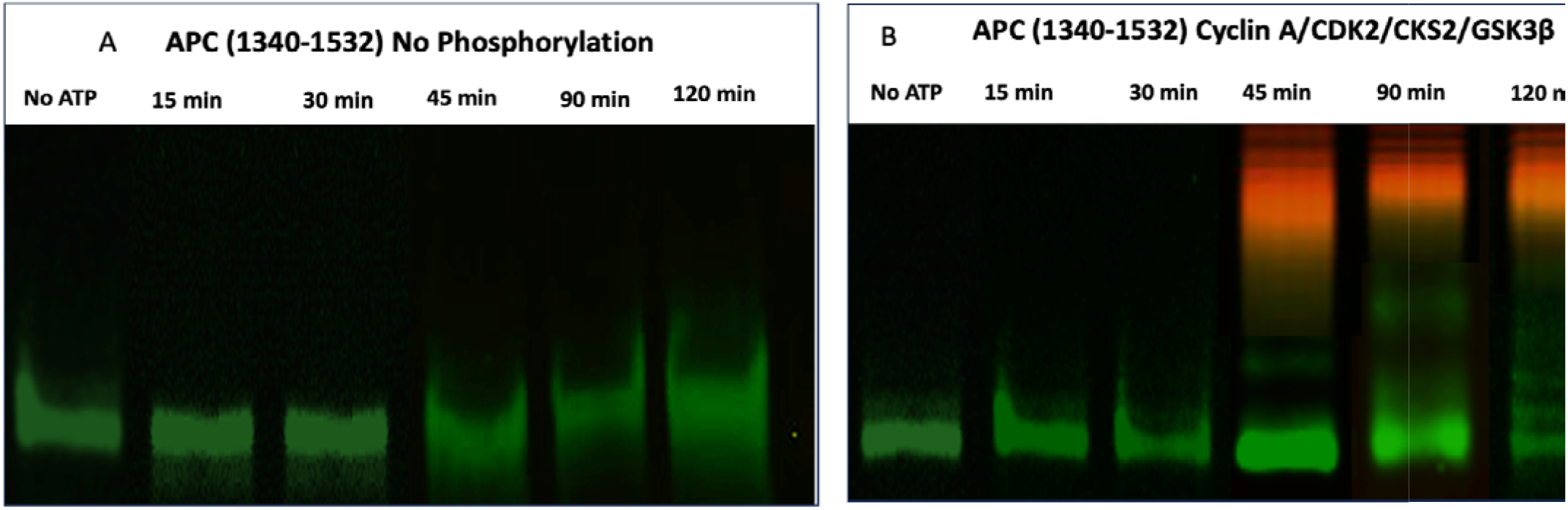
Rate of β-catenin ubiquitination occurs faster when APC CID undergoes in vitro phosphorylation with Cyclin A/Cdk2/Cks2/ GSK3β. In vitro ubiquitin transfer assay. Fluorescent image of SDS PAGE with trace fluorescently labeled ubiquitin (red) and β-catenin (green). These fluorescent proteins were incubated for the indicated times with UBA1, Ubc5h, and SCF^BTRC^ in the presence of APC CID with either A) no phosphorylation or B) phosphorylated with Cyclin A/Cdk2/Cks2/ GSK3β

### Phosphorylation of the CID regulates β-catenin signaling

When β-catenin is stabilized by Wnt signaling, it goes into the nucleus and binds TCF transcription factors, acting as a transcriptional co-activator. The role in Wnt signaling of CDK and GSK-3 phosphorylation-dependent binding of APC’s R2-CID region to β-TrCP1 was tested using the standard β-catenin gene regulation assay (TOPFlash) that links luciferase to a multimerized TCF promoter^53^. Increased TOPFlash activity indicates stabilization of β-catenin, i.e., loss of destruction activity.

We employed APC4 cells, which are truncated in both alleles by CRISPR to before the start of the R2 region^35^. Previous work revealed that expression of GFP-fused R2-SAMP1 can downregulate TOPFLASH activity in this cell line. To assess the importance of phosphorylation of this region, we tested point mutations in the hypothesized CID pSGxxpS motif that either disrupted phosphorylation or enabled priming of GSK-3 to constitutively phosphorylate of the CID (Figure 5). APC4 cells were transfected with endogenous R2-R3 where Ala or Glu (phosphomimetic) replacing the Ser residues at 1411, 1415, or 1419. We also tested the conserved Gly at position 1416. Mutating S1411 had little effect, indicating that either the first potential β-TrCP1 binding site 1410-1415 is not important, or that the neighboring site 1414-1419 suffices for β-TrCP1 binding and β-catenin destruction. In contrast, changing S1415 to Ala, which would eliminate priming of S1411 and eliminate the key pSer of the second possible β-TrCP1 binding site, increased TOPFlash activity about 5-fold. Changing S1415 to Glu had little effect; the small increase in TOPFlash activity could arise from Glu not serving as a perfect mimic of pSer at this position. Disabling GSK-3 priming with the S1419A mutation elevated TOPFlash signal comparable to S1415A. Constitutively priming the GSK-3 sites with the S1419E mutation gave an intermediate TOPFlash signal, which may indicate that the Glu does not precisely mimic pSer in this position for GSK3 priming. Furthermore, mutation of the conserved Gly at 1416 to Ala yielded the largest increase in TOPFlash activity, indicating a large disruption of β-catenin destruction. Overall, the TOPFlash data are consistent with a dual kinase phosphorylation of the CID generating a β-TrCP1 binding site that is required for efficient β-catenin destruction.

To determine if binding was significantly affected by these mutations APC tagged with GFP was immunoprecipitated with anti-GFP nanobody resin and blotted against β-TrCP and the GFP tag (Figure 5b). This showed that the APC construct could readily bind to and Co-IP β-TrCP and the mutations had a negligible effect on total binding (Figure 5c). To determine if our hypothesis that APC was stabilizing β-TrCP, we blotted the whole cell lysate for β-TrCP with and without APC transfection (Figure 5d). Finally, to demonstrate this region of APC was sufficient to rescue destruction complex activity we treated APC4 cells with the proteasomal inhibitor MG132 and blotted the lysate against β-catenin (Figure 5e). This showed rescue of ubiquitination of β-catenin to corroborate the TOPFLASH data.

## DISCUSSION

The E3 ubiquitin ligase SCF^β-TrCP1^ is responsible for the ubiquitylation of phosphorylated β-catenin, but its association with and regulation by the β-catenin destruction complex has been challenging to characterize. Some reports have indicated that APC associates with SCF^β-TrCP1^ ^38^, but the mechanisms mediating and regulating this interaction remain unclear. We have explored the role of the highly conserved sequence between APC repeats 1 and 3, the R2 and CID regions. It is deleted in many cancers and genetic studies reveal it is essential for APC function^23,33,54,55^.

We used a mass spectrometry screen to identify binding partners of the R2/CID region. This revealed that cyclin-dependent kinases associated with the R2-CID region of APC. Based on the conserved consensus GSK-3 and CDK substrate recognition motifs, we hypothesized that these enzymes phosphorylate the R2-CID, which would then mediate binding to β-TrCP1. We tested this hypothesis using multiple biochemical approaches, and this revealed that APC R2/CID phosphorylation elevated binding to SCF^β-TrCP1^ and enhanced the rate of ubiquitin transfer to β-catenin *in vitro*. This defines the mechanisms explaining the crucial role for the APC CID in β-catenin destruction.

The affinity of the Cdk2/GSK3 phosphorylated APC CID binding to β-TrCP1, compared to the interaction with β-catenin, provides a plausible mechanism for an essential role for APC in the regulation of β-catenin destruction. The faster on and off rates, which lead to the 15x higher affinity for β-catenin, and the faster dissociation than association rate with phosphorylated-CID means that β-catenin will be the preferred β-TrCP1 partner (with caveat that the actual concentration of β-TrCP1 in the complex is not known) in the presence of both binding partners. However, if β-catenin dissociates (i.e., during active destruction), binding to the R2-CID region will stabilize β-TrCP1 in the destruction complex rather than allow it to dissociate from the APC region. Furthermore, the low on-rate of APC binding to β-TrCP1 supports data showing APC cannot, by itself, recruit β-TrCP1 to the destruction complex, only retain it. This stabilizing behavior provides a kinetic trap where the CID effectively acts as a scaffold by retaining β-TrCP1 in proximity to the complex.

Our mass spectrometry data revealed the presence of β-TrCP1, CAND1, and other components of the SCF^β-TrCP1^ complex, associate with the APC R2-CID region. The E3 ligase SCF^β-TrCP1^ is composed of the scaffold Cul1, the RING domain protein Rbx/Roc1, the adaptor Skp1, and the substrate-binding F-box protein β-TrCP1^56,57^. Neddylation of the Cul1 protein promotes a conformation that efficiently transfers ubiquitin from the E3 to lysines on the F-box–bound target^58–60^ and prevents disassembly of the E3 complex. There are 69 different F-box substrate receptor proteins encoded in the human genome, the majority of which have been detected, implying that there are many SCF complexes^61^. CAND1 mediates rapid exchange of different F-box proteins from the SCF, which enables fine-tuned responses to changing levels of SCF substrates. In the absence of a ubiquitylation substrate, the COP9 signalosome (CSN) can de-neddylate the SCF, which favors CAND1-mediated exchange of F-box partners. However, the presence of a substrate bound to an F-box protein stabilizes SCF^Fbox^ complexes, including β-TrCP, by inhibiting de-neddylation of the SCF by the CSN and thereby preventing exchange with other F-box proteins^62–66^.

We propose that the APC CID inhibits, when appropriately phosphorylated, CAND1 mediated SCF exchange by providing a β-TrCP substrate mimic that stabilizes SCF^β-TrCP^ when β-catenin is not present or is undergoing proteasomal degradation. Relative to true phosphoserine degrons, this mimic binds differently with slower on and off kinetics and serves to stabilize and retain this “pseudo substrate” to β-TrCP to ensure that the destruction complex is active. The ubiquitylation itself is processive and rapid^51^, and we believe that the CAND1 exchange of β-TrCP that has dissociated from ubiquitylated intermediates is not significant. As the phosphorylated and ubiquitylated β-catenin is removed by the proteasome, the phosphorylated CID will bind β-TrCP1 and thereby prevent F-box substrate receptor exchange by the actions of the CSN and CAND1. Although the kinetics of β-catenin removal and rebinding of another β-catenin substrate has not been measured in cells, studies of model substrates *in vitro* have shown that the processes of poly-ubiquitin binding, substrate binding, de-ubiquitylation, substrate unfolding, and destruction occur over multiple seconds to minutes^67^. In contrast, CAND1 mediates rapid F-box exchange on the order of a second. Thus, it is probable that in the absence of the preferred CID phosphorylation, β-TrCP1 would be exchanged during the destruction of β-catenin and thereby lower the overall efficiency of β-catenin destruction. Stabilizing β-TrCP1 by binding to the phosphorylated CID favors retention of the SCF^β-TrCP1^ in the destruction complex and ensures that the SCF^β-TrCP1^ remains available for β-catenin ubiquitylation. Thus, observations argue that the essential role of the APC CID is to increase the efficiency of ubiquitin transfer, which would otherwise be too slow to prevent target gene activation by β-catenin. This would explain the oncogenic effect of APC truncation mutants lacking the CID.

Earlier observations are consistent with our proposed model. Li and colleagues found reduced amounts of β-TrCP1 in immunoprecipitation experiments from cells expressing APC lacking just its CID^35^. Ranes et al described an elegant reconstitution of the destruction complex using recombinant proteins^38^. They found that full length APC or large fragments of APC, along with Axin and the other components of the destruction complex including the GSK-3 and CK1, enhanced ubiquitylation of β-catenin. Unlike our data, however, this did not require the CID. However, we found that CDKs, rather than CK1, are needed to prime the required GSK-3 phosphorylation of the APC CID. Moreover, even with the correct (or adventitious) priming, the concentration of APC used in their ubiquitin transfer assays were too low to observe the effect of the phosphorylated APC CID on β-TrCP1 activity. Nonetheless, why the data in this study showed enhancement of ubiquitin transfer activity could be due to several possibilities. As we noted above, we did not include Axin in our experiments. It is possible that Axin might effectively increase the local concentration of β-catenin to enable some enhancement of ubiquitin transfer; the concentrations of GSK-3 and CK1 in their assays were high and this would result in higher affinity binding^24,26^. Ranes et al. and others^7,68,69^ have found that fragments of APC substantially longer than the minimal R2-R3 sequence used in our transfer assays promote destruction of β-catenin. It is possible that regions of APC outside the R2-R3 contribute to β-TrCP1 binding^38,70^ and might act as another potential scaffold to enhance ubiquitin transfer or other interactions with the SCF complex that contribute to enhanced ubiquitin ligase efficiency independent of the effect of the phosphorylated CID proposed here. We have not been able to produce a biochemically stable, homogenously phosphorylated, longer construct of APC to test this hypothesis. Nonetheless, our results clearly show an effect of CDK priming and GSK-3 phosphorylation of the APC CID on the rate ubiquitin transfers to β-catenin.

Our model (Figure 7) predicts that loss of the CID reduces the efficiency of β-catenin destruction by allowing rapid exchange with other F-box proteins, such that destruction activity falls below a threshold needed for appropriate regulation of β-catenin levels. Quantitatively assessing the effect of this stabilization is challenging given the many unknowns in the system, including its dynamic stoichiometry^38^ and the possibility that the destruction complex is a phase separated species^28^. Also, whereas R2 and the CID are conserved and likely have unique roles, R3 does not have a privileged mechanistic role in β-catenin destruction, although it is the most strongly binding 20mer repeat^26^. Deletion of various 20mer repeats showed that destruction will work if there is at least one 20mer present^23,32^. The deletions retain at least one 20mer near the R2-CID region, consistent with a scaffolding role of APC maintaining the β-catenin substrate in proximity to the E3. Nonetheless, the differential binding affinities of different repeats could have a role in modulating the residence time of β-catenin and also affect destruction complex turnover^23,24^. In the multivalent destruction complex, the multiple repeats help to retain β-catenin, which likely increase the effective concentrations of β-catenin and therefore affect the kinetics of binding in the proximity of β-TrCP1.

**Figure 7).**
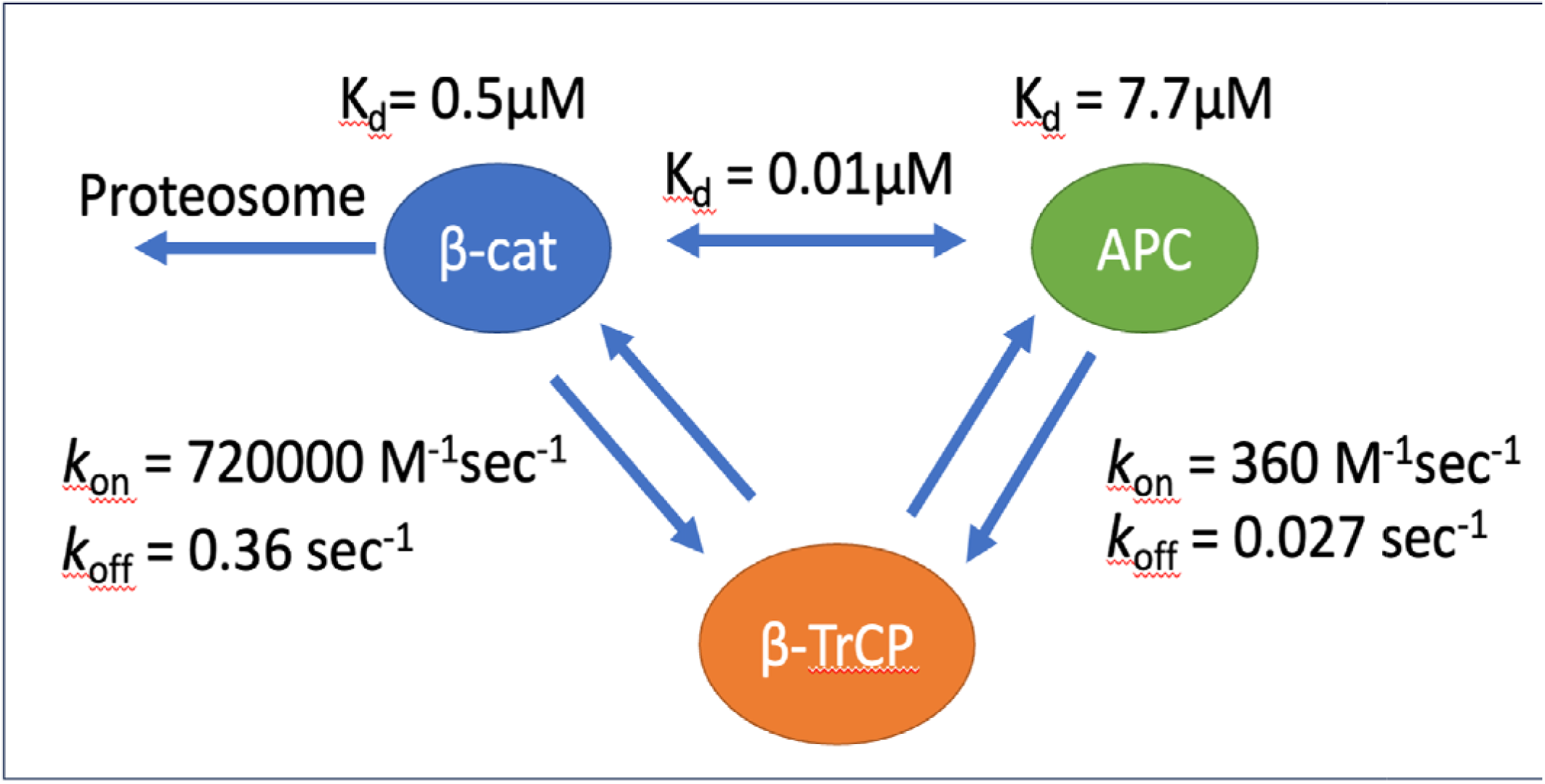
Kinetic model of APC as it interacts with BTRC and Beta-Catenin

In this study we also observed that the Cyclin/Cdk complexes associated with the APC CID region. Previous studies have shown that Cyclin E, which recognizes the same binding interface as Cyclins A and B^71–74^, requires GSK3 activity for its proteasomal destruction after self-priming by Cdk2^75^. Since our model places the Cyclin/Cdk complex in close proximity to GSK3, it is likely that the R2/CID region participates in the proteasomal degradation of CyclinE, at a minimum in scaffolding the phosphorylation of the two CyclinE degrons by GSK3^75,76^. As noted above, CAND1 rapidly exchanges E3 substrate receptors and this has been demonstrated between β-TrCP and FBXW7^63^, the E3 responsible for Cyclin E ubiquitination^76^. While beyond the scope of this work, a consensus motif in the APC CID region mirrors the CyclinE degron and could perform a similar function with FBXW7 as presented here with BTRC. This would allow Wnt activation to directly affect cell cycle progression as inhibition of GSK3 would stabilize the pool of CyclinE bound to APC in addition to halting Beta-catenin degradation. This provides clear direction for future studies.

We showed that the conserved APC CID regulates ubiquitylation of β-catenin in a CDK/GSK-3 dependent manner. Our findings provide a mechanism for the essential role of the CID. The unexpected role of cell cycle kinases in this process controlling SCF activity through APC has a more general role in regulating the cell cycle. This connection, although beyond the scope of this study, will be needed to understand the role of APC in cellular growth. The prospect of multiple substrates that could regulate the ubiquitylation and destruction of cell cycle components could potentially impact not only APC biology but more general aspects the cell cycle, including the timing of the cycle and how different CDKs can contribute to specifically timed destruction of different components represents a key area to investigate.

## Methods

### Cloning constructs

pEBB-HA-SBP, his-Ubch5, and his-UBA1 were acquired from Addgene. The His-Cdc34b vector was a gift from Dr. Brenda Schulman. SMT3-BTRC/Skp1 and Avi-SMT3-BTRC/Skp1 in pCDF-dual were a generous donation from Nurix Inc. The USP2cc-his vector was a gift from Dr. Ron Kopito. Vectors for Cdk2/CyclinA/Cks2 were a gift from Dr. David Barford. APC constructs were cloned into pGEX-TEV-Trp or pGEX-TEV. APC phosphomimics were inserted by site-directed-mutagenesis (QuickChangeII Agilent).

### Protein Purification

CyclinA/Cdk2/Cks2 complex was purified per publication^77^

SMT3-BTRC/Skp1 and Avi-SMT3-BTRC/Skp1 was purified per publication ^45^ USP2cc, UBA1, Cdc34b, ULP1, TeV, GSK-3b, Ck1a were purified per (supplemental table 3)

### Purification of cyclin-Cdk-Cks complexes

Human cyclin-Cdk and cyclin-Cdk-Cks fusion complexes were purified from budding yeast cells using a FLAG affinity purification method, modified from a previous protocol used for HA-tag purification^78^. Briefly, N-terminally 3XFLAG-tagged cyclin-Cdk and cyclin-Cdk-Cks fusions were cloned into high-copy yeast vectors using a glycine-serine linker and overexpressed from the GAL1 promoter. The overexpressed 3FLAG-tagged complexes were then purified by one-step immunoaffinity chromatography using ANTI-FLAG M2 affinity agarose beads (Sigma-Aldrich A2220) and eluted with 0.2 mg/mL 3XFLAG peptide (Sigma-Aldrich F4799)^79^.

### Purification and *in vitro* phosphorylation of GST-APC

GST-APC constructs were purified as follows. 20mL of an overnight culture was inoculated into 2L of autoclaved LB media. This was grown to an OD600 of 1.2-1.5 at 37C and induced for 2hour at 37C by the addition of 1mM IPTG. The cultures were centrifuged at 8000rcf for 15min, resuspended by vortexing in 30mL of 20mM Tris pH 8.0 300mM NaCl 10mM DTT, and frozen at-80C. Pellets were defrosted and lysed by two passes of an emulsiflex at 15,000psi after the addition of protease inhibitors and DNAse1. Post-lysis, 2mM EDTA and fresh PMSF was added. Lysate was clarified by ultracentrifugation at 41,000rcf for 30min. Clarified lysate was applied to 10mL of freshly cleaned GST-agarose resin and batch bound at 4C for 30min with gentle rotation. Resin was washed with 10CV of 1M NaCl 20mM Tris pH 8.0 10mM DTT, followed by 10CV of lysis buffer. Protein was eluted in 3 CV of lysis buffer supplemented with 10mM reduced glutathione. Protein was diluted 1:4 in MonoQ running buffer A (20mM Tris pH 8.0 10mM DTT) and subjected to anion-exchange chromatography. Fractions containing APC were pooled and re-bound to 10mL of freshly cleaned GST-agarose and washed with phosphorylation buffer 20mM Tris pH 8.0 300mM NaCl, 5% glycerol, 1mM DTT, supplemented with 10mM MnCl2. Lambda phosphatase was added, and resin was incubated overnight at 4C.

### *In vitro* phosphorylation of the APC protein

Resin was washed first with 5CV phosphorylation buffer supplemented with 10mM EDTA, followed by 5CV of phosphorylation buffer supplemented with 10mM MgCl2, before addition of 2mM ATP pH 8.0, protease inhibitors and ∼500nM of kinases. (Cdk2/Ck1a/GSK-3b) Resin was incubated for 2hours at 30C. Protein was eluted in MonoQ Urea Buffer A supplemented with 10mM Glutathione. (4M Urea, 20mM Tris pH 8.5, 10mM DTT) and subjected to a very shallow MonoQ gradient^24^

Proteins were analyzed for phosphorylation by Diamond Pro-Q phosphostain and a shift on the MonoQ chromatogram. Phosphorylated material was concentrated to 20uM by 10kDa spin concentrator at 4000rcf in 5min intervals and then buffer exchanged via SEC (agS75 or agS200). Protein was aliquoted and flash frozen in liquid nitrogen and stored at-80C.

### In vitro kinase assays

*In vitro* kinase assays were performed as described^80^. Briefly, separate reactions contained an equal amount of substrate (in the range of low uM) and equal amount of purified kinase complexes (in the range of low nM). Aliquots were collected at two time points (8 and 16 minutes) and the reaction was stopped with SDS-PAGE sample buffer. The basal composition of the *in vitro* kinase assay mixture contained 50 mM HEPES pH 7.4, 150 mM NaCl, 5 mM MgCl_2_, 0.02 mg/ml 3XFLAG peptide, 5% glycerol, 3 mM EGTA, 0.2 mg/ml BSA and 500 μM ATP with 2 μCi of [γ-32P] ATP added per reaction (PerkinElmer). Histone H1 protein used as a general substrate for Cdk was purchased from EMD Millipore (14-155). Phosphorylated proteins were separated on 10% SDS-PAGE gels. Phosphorylation of substrate proteins was visualized using autoradiography (Typhoon NIR Plus; Cytiva). Autoradiographs were visualized and quantified with the ImageQuant Software (Cytiva).

### Coelution

Purified APC CID 1300-1586 (70uM) phosphorylated with CyclinA/Cdk2/Cks2 + GSK3β, CK1 + GSK3β, or non-phosphorylated APC was mixed with β-TrCP1/Skp1 (30uM) in a 1.5 mL tube and incubated on ice for 1 hour. Each of these samples were ran over a S200 analytical column at 0.5 mL/min. Peaks were noted on chromatographs and fractions were analyzed by SDS-PAGE.

### Bio-Layer Interferometry (BLI)

β-TrCP1/Skp1 was avi-tagged and biotinylated using NHS-PEG4-biotin (ThermoFisher). BLI measurements were performed with Octet Red384 (Sartorius) and a GatorPrime (Gator Bio) using streptavidin biosensors. All steps were performed at 25°C shaking at 1000 rpm. Data sets were collected at 10 Hz. Each streptavidin biosensor tip was hydrated in 200 uL of the same SEC buffer (20 mM Tris pH 7.6, 150 mM NaCl, and 1 mM EDTA) for 10 minutes. We tested each individual protein for non-specific interactions with the biosensors. Next, we added 0.1 uM BSA and 0.01% Triton X to the SEC buffer to reduce non-specific interactions and decrease background noise. APC CID samples were diluted to concentrations ranging from 20 to 0.625 uM in binding buffer (20 mM Tris pH 7.6, 75 mM NaCl, 1 mM EDTA with 0.1 uM BSA and 0.01% Triton X). Biotinylated β-TrCP1/Skp1 was diluted using the same binding buffer to a concentration of 50 uM and loaded on streptavidin biosensors for 2 minutes. Sensors were re-equilibrated in binding buffer for 15 minutes, and then placed in wells containing binding buffer for 1 minute to establish a baseline. Biosensors were then placed in 90 uL of APC CID ligand at varying concentrations to monitor the signal for association. Next, each biosensor was place in a well containing only the binding buffer to monitor dissociation. The last well contained binding buffer without ligand to define baseline drift, and experiments were performed in triplicate.

We us the Octet Data Analysis 10.0 or Gator 1.7 software to process all data initially. Biosensor number 8, the sensor placed in the last well containing binding buffer without ligand, was used as the reference sensor and its signal was subtracted from the signals obtained by other biosensors. Corrected binding signals were exported to Prism 9 GraphPad for curve fitting, association-dissociation kinetic data and a one-site binding model. The wavelength shifts recorded at 375 seconds after the start of association were graphed as a function of the corresponding APC CID concentration to calculate the K_D_ values. The K_D_ and coefficient of determination (R^2^) values were calculated and shown on the graph. Each assay was performed in triplicate.

### Ubiquitin transfer assays

Ubiquitin and β-catenin were trace fluorescently labeled with Texas Red and Carboxyrhodamine (respectively) as previously described^51^. E1 UBA1 (50nM or 1uM), E2 Cdc34b (1uM or 4uM), fluorescently labeled Ub (60uM) were mixed separately in 1X Transfer buffer. (20mM Tris pH 7.6, 100mM NaCl, 1mM DTT, 10mM MgCl2). APC, β-catenin and APC were mixed in that order into 1X transfer buffer. Reaction was initiated by addition of 2mM ATP pH 7.6 to the E1 containing tube, incubated for 5min at RT. E1 mixture was then added into E3 mixture to initiate Ub transfer to β-catenin and samples were taken from the common tube at defined timepoints and quenched by addition of SDS-Loading buffer containing 100mM BME.

### Cell Culture

HEK 293 cells were grown in DMEM 10% FBS supplemented with gluatamine and penicillin/streptomycin. C59 and CHIR99021 were dissolved in DMSO at 100uM per mfc recommendations.

### Co-immunoprecipitations

HEK 293 cells with a CRISPR truncated APC (4-3 line), a kind gift from Vivian Li, were grown to 80% confluency in 7.5cm plates. Cells were transfected with GFP-APC mutants with Lipofectamine-2000 and allowed to grow for 48 hours.

Cells were harvested by trituration in NP-40 Buffer (1% NP-40, PBS, 10% Glycerol, 10mM beta-glycerophosphate, 5mM pyrophosphate, protease inhibitors, 2mM EDTA, 5mM DTT). Lysate was incubated on ice for 10 minutes and then centrifuged at 10,000 rcf for 10min. The supernatant was applied to 20uL of GFP-nanobody resin and incubated for 30min at 4C on a small rotator. Beads were washed three times with lysis buffer and eluted by the addition of 50uL of 100mM Glycine pH 2.7 100mM NaCl. Sampled were quenched by the addition of 1M Tris pH 8.5 and then run on SDS-Page 200V for 40min. Samples were transferred to nitrocellulose and blocked in 5% milk TBST before being blotted with primary antibody. Following washes and secondary staining, blots were read on a Licor infrared scanner.

### TOPFlash Reporter Assays

HEK 293 cells with a truncated APC were seeded at 10,000 cells/well into an opaque 96 well plate and allowed to grow overnight. Cells were then transfected with Super-TOPFlash, LacZ, and APC-GFP vectors and allowed to grow for 48 hours. Cells were then subjected to the Dual-light reporter kit and read out in a luminometer. Plates were incubated for 1 hour on a shaker and then LacZ signal was assayed per mfc instructions.

### Cell Growth Assays

HEK 293 cells with a truncated APC were grown to 80% confluency in 10cm plates. Cells were transfected with APC-GFP vectors and lipofectamine 2000 and incubated overnight. Cells were then trypsinized for 5min, quenched in 10mL of DMEM, counted and seeded at 500,000 cells per 5 cm flask and allowed to grow for 48 hours. Cell flasks were then trypsinized for 5min, quenched in 10mL of DMEM, counted by coulter counter.

### Mass Spectrometry

#### Preparation of Mass Spectrometry Samples

293T cells were grown in DMEM with 10% FBS deficient in Leucine and Methionine in five 10cm plates per condition for 24hours. Cells were transfected with pEBB-SBP-R2/CID-HA or pEBB-SBP-R2/R3-HA using 60uL of Lipofectamine 2000 and 20ug of DNA per plate and allowed to recover overnight. Media was replaced with DMEM with 10% supplemented with 2mM Photo-Methionine and 4mM Photo-Leucine (Pierce) and allowed to grow for 24hr. Cells were treated with 10uM of MG132(Selleckchem) in DMSO for 2 hours prior to harvesting.

Cells transfected were washed twice with 37C DPBS. Cross-linking was activated by exposure to 365nm UV light at 0.5cm for 25min.

#### Mass Spectroscopy Samples were then prepared in two ways

Lysis Buffer = 20mM HEPES 7.5 150mM NaCl 0.1mM EDTA and either 0.2% NP-40 (Gel Sample Preparation) or 0.6% Triton X-100 (Whole Cell Preparation)

#### Whole cell lyophilized preparation

Cells were harvested by cell scraping in 1.5mL of lysis buffer per plate and lysed by trituration. Lysis was confirmed by performing a trypan blue staining on 2uL of material and visualizing on a microscope. Cells were pelleted at 21,000 rcf for 10min. Clarified lysate was incubated with 50uL of Streptavidin resin per plate for 2 hours with rotation. Resin was washed three times with 1mL of lysis buffer lacking detergent. Samples were eluted with 2mM d-biotin in 150uL of 20mM HEPES 7.5 150mM NaCl. Samples were pooled, then lyophilized and submitted for LC-MS/MS.

#### SDS PAGE Gel samples preparation

Cells were harvested by cell scraping in 1.5mL of lysis buffer per plate and lysed by trituration. Lysis was confirmed by performing a trypan blue staining on 2uL of material and visualizing on a microscope. Samples were spun at 1000rcf for 5min. The plasma membrane fraction was removed by pipetting off the top 100uL of the supernatant. Cells were then spun at 21,000rcf for 30min. Supernatant was removed and retained for analysis. The pellets were pooled and dissolved in 750uL of RIPA buffer and incubated on ice for 10min. The solubilized material was placed on an 4mL 80/75/55/35% percoll gradient in 20mM HEPES 7.5 150mM NaCl 0.1mM EDTA 0.6% Triton X-100 and spun for 2 hours at 1000rcf. 10uL of each 400uL fraction, the plasma membrane, and the lysate supernatant were analyzed by western blot against the HA tag to identify fractions containing cross-linked material. Material was found in fractions 3-7 of the percoll gradient. Percoll was removed from the positive fractions by ultracentrifugation at 100,000 rcf for 90min. The supernatant was incubated with 20uL of Streptavidin resin for 2 hours with rotation. Resin was washed three times in 1mL of lysis buffer lacking detergent. Resin was eluted in 50uL of 20mM HEPES 7.5 150mM NaCl containing 2mM d-biotin for 1 hour. Eluted material was run in SDS-PAGE and bands were extracted based on a mass spec compatible silver stain kit (ThermoFisher-Pierce). Silver stain bands correlating to western blot sizes were excised from the gel in a sterile hood. Gel pieces were treated to remove silver as per mfc instructions and washed three times in sterile deionized water before being submitted for LC-MS/MS analysis.

#### Mass spectroscopy Instrument Run and Sample Prep

Gel pieces were cut and resuspended in 50mM Ammonium Bicarbonate and 5mM DTT and heated at 55C for 30min. Tubes were allowed to cool and dried in a speed vac before resuspension in 50mM Ammonium bicarbonate, 5 pmol Promega sequencing grade trypsin and 0.02% Promega protease max. Tubes were incubated for 2 hours at 50C. Protein was extracted with 1% formic acid, 66% acetonitrile 33% 100 mM Ammonium bicarbonate, incubated for 10min at 37C and spun down at 10,000 rcf for 2min and stored at-4C.

#### Mass Spectroscopy

In the mass spectrometry experiment, dried samples were resuspended in 2% aqueous acetonitrile with 0.1% formic acid and injected onto either an LTQ Orbitrap Velos hybrid mass spectrometer (Thermo Scientific, San Jose, CA) with liquid chromatography using a nanoAcquity UPLC (Waters Corporation, Milford, MA) or Orbitrap Fusion Tribrid mass spectrometer (Thermo Scientific, San Jose, CA) with liquid chromatography using an Acquity M-Class UPLC (Waters Corporation, Milford, MA). For liquid chromatography, a flow rate of 450 nL/min was used, where mobile phase A was 0.2% formic acid in water and mobile phase B was 0.2% formic acid in acetonitrile. Analytical columns were pulled and packed in-house using 360 micron fused silica with an I.D. of 100 microns packed with Dr. Maisch 1.8 micron C18 stationary phase to a length of ∼20 cm. Peptides were directly injected onto the analytical column using a gradient (2-45% B, followed by a high-B wash) of 90 min.

The mass spectrometer was operated in a data-dependent fashion using CID or HCD fragmentation for MS/MS spectra generation collected in the ion trap.

## Data Analysis

The collected mass spectra were analyzed using both SEQUEST in Scaffold (Proteome Software, Portland, OR) and Byonic (Protein Metrics, Cupertino, CA) for peptide identification and protein inference. The search was performed against a targeted list of proteins based on the assay. For cross-linked peptides, this database of proteins was expanded to include additional known false positives to better set thresholds for crosslink assignments. Cysteine modified with propionamide was set as a fixed modification in the search, with other post-translational modifications, e.g. oxidation of methionine, included as variable modifications. Where appropriate, cross links were investigated as described above using the Byonic X-Link node to define linked peptides, and validated by inspection in Byologic (Protein Metrics, Cupertino, CA). Data were held to 6 ppm mass tolerance for precursors and 0.4 Da for MS/MS fragments in Byonic, allowing up to two missed cleavage sites. Data were validated using the standard reverse-decoy technique at a 1% false discovery rate.

Data for SEQUEST searches were held to a 3ppm mass tolerance for precursors and gated peptide spectra with a Xcorr >2.5.

## Acknowledgements

We would like to thank Vivian Li for the APC4 cells. We would like to thank David Barford, Brenda Schulman, and Ron Kopito for expression plasmids. We also appreciate the efforts of Elise Bruguera and Lenore Urbani for early contributions to this study. The authors would like to thank James Ferrell and Ron Kopito for critical feedback. The authors would like to thank Mark Peifer and Roel Nusse for a critical reading and valuable insights on the manuscript.

Mass spectrometry measurements were performed at the Vincent Coates Foundation Mass Spectrometry Laboratory, Stanford University Mass Spectrometry (SUMS - RRID:SCR_017801).

This work was supported in part by NIH P30 CA124435 utilizing the Stanford Cancer Institute Proteomics/Mass Spectrometry Shared Resource. This work utilized the LTQ-Orbitrap mass spectrometer system (RRID:SCR_018694) that was purchased with funding from National Institutes of Health Shared Instrumentation Grant S10RR027425. Additional support to Darius Johnston from NIH NRSA 5F31CA196170-02 and to William Weis through NIH R35 GM131747

## Author contributions

DJ and WW conceived of the project. DJ and CLT designed and performed biochemical and biophysical experiments. MK performed radioactive kinase assays. DJ, LF, RL, and CA performed mass spectrometry experiments and analysis. DJ, CLT and WW analyzed data. DJ prepared the manuscript. DJ, CLT and WW edited and finalized the manuscript.

