## Supplemental Figures for "Phosphorylation-dependent binding of the Adenomatous Polyposis Coli protein to β-TrCP1 regulates β-catenin destruction"

Figure S1

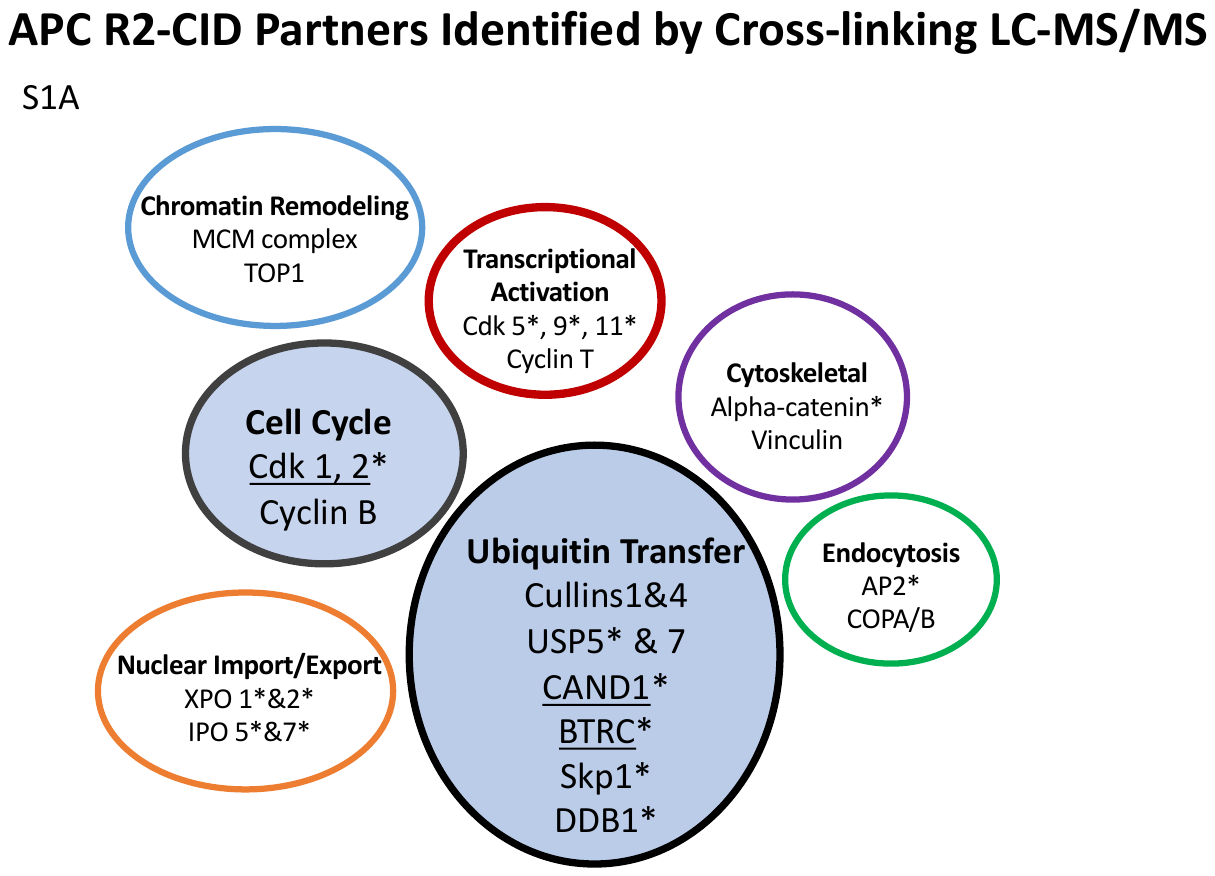

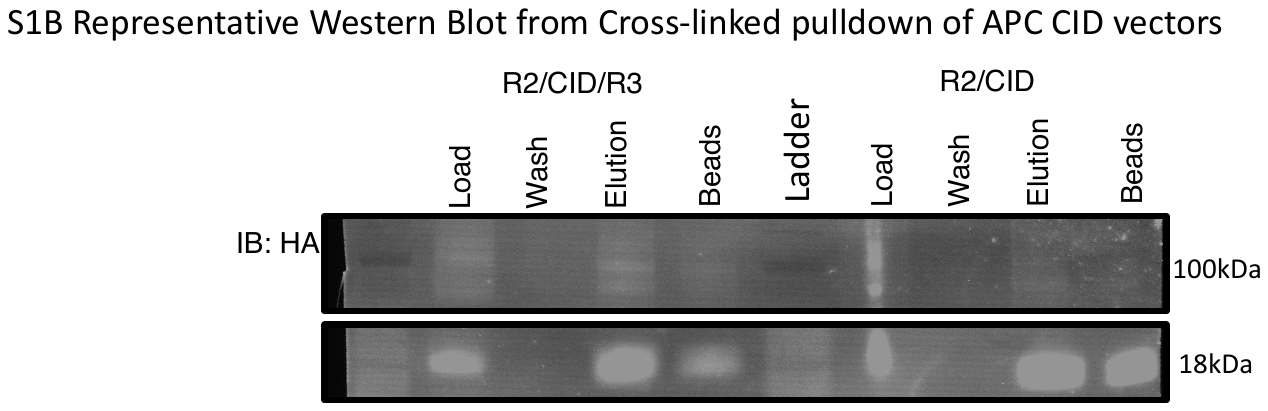

Figure S1A: The APC CID region interacts with Ubiquitin Transfer Components and Cdk’s A summary of MS Hits from the cross-linking mass spectroscopy screen. *Denotes a cross-linked hit in the samples.

B: Representative cross-linking anti-HA Western Blot showing APC “bait” shifting upwards in size

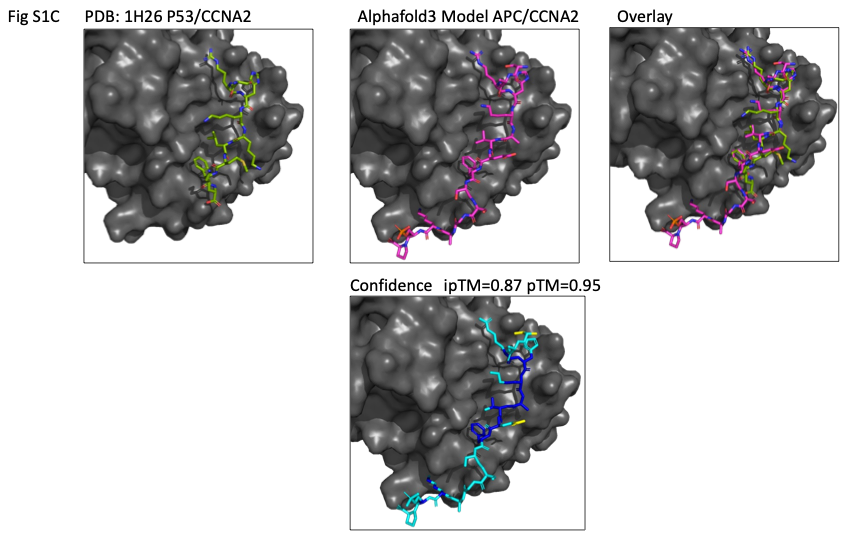

Figure S1C) Alphafold3 modeling of the predicted APC/CyclinA interface. Using PDB:1H26 (CyclinA/P53) as a template, confidence for the model is pTM=0.87 and pTM=0.95

Table S1 and S2
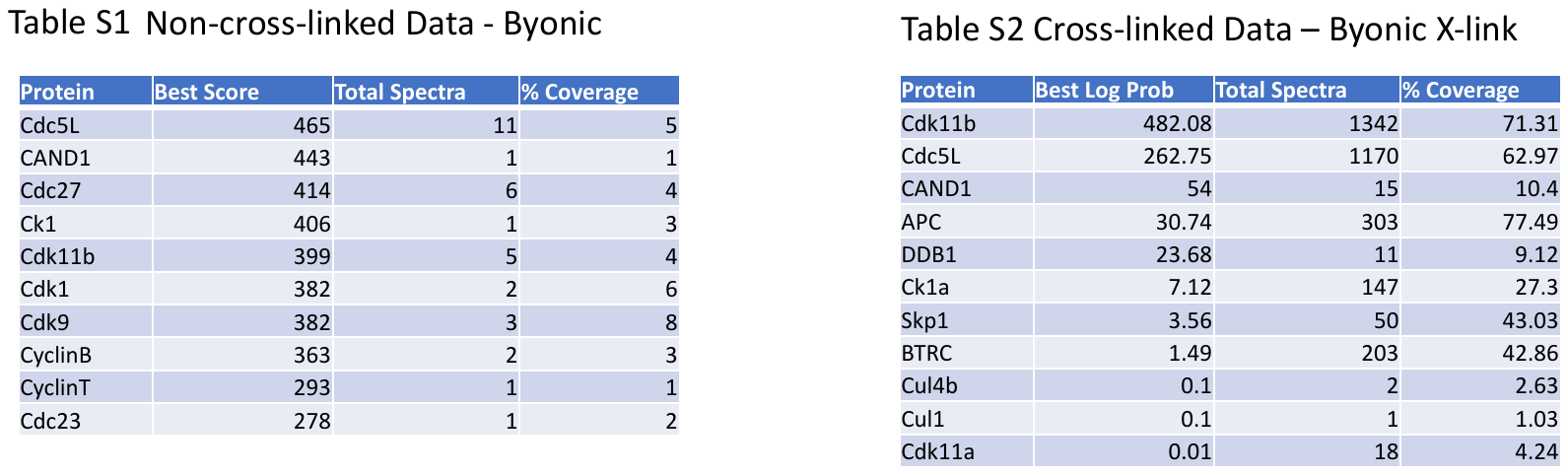

Table S1: Non-cross-linked LC-MS/MS data analyzed in Byonic.

Table S2: Cross-linked LC-MS/MS data analyzed in Byonic X-link.

| Supplemental Table S3 |  |  |  |  |  |  |
| --- | --- | --- | --- | --- | --- | --- |
| Protein | Construct | Strain | Expression System | Purification | Comment | Reference |
| Beta-catenin | pET-GST | BL21(DE3)  RIL | E.coli | GST, Tev Digest, MonoQ,  S200 | - | ^24^ |
| APC-GST | pET-W-GST | BL21(DE3)  RIL | E.coli | See Methods | - | This Study |
| APC HA-R2-CID-SBP | pEBB-SBP-HA | HEK293 | Mammalian | Cell Fractionation  Streptavidin |  | This Study |
| APC HA-R2-R3-SBP | pEBB-SBP-HA | HEK293 | Mammalian | Cell Fractionation  Streptavidin |  | This Study |
| BTRC1/Skp1 | pFAST-BAC | Sf9 | Insect | GST, Thrombin digest, MonoS, S200 | Gift from Brenda Schulman | (Li et al., 2005) |
| BTRC1/Skp1  delN 139-569 | pCDF-Duet1 | Lemo21(DE3) | E.coli | NiNTA, ULP1 digest, Reverse NiNTA, MonoS, S200 | Gift from Nurix  Co-expression | ^45^ |
| BTRC1/Skp1  delN 139-569-avi | pCDF-Duet1 | Lemo21(DE3) | E.coli | NiNTA, ULP1 digest, Reverse NiNTA, MonoS, S200 | Gift from Nurix  Co-expression | ^45^ |
| ULP1 | pET28b | BL21(DE3)  RIL | E.coli | NiNTA, S200 |  |  |
| Cul1 NTD Rbx1-Cul1 CTD | pAL Cul1 NTD, pCool Rbx1 Cul1 CTD | BL21(DE3)  RIL | E.coli | GST, Thrombin Digest, MonoS, S200 | Gift from Brenda Schulman | (Li et al., 2005) |
| Cdk1/CyclinB/Cks1 | pRS425-pGAL1 | DOM0071  (Gift from David Morgan) | Yeast | 3FLAG, M2 affinity | Gift from Skotheim Lab | Topacio et al., 2019 |
| Cdk1/CyclinB/Cks2 | pRS425-pGAL1 | DOM0071  (Gift from David Morgan) | Yeast | 3FLAG M2 affinity | Gift from Skotheim Lab | Topacio et al., 2019 |
| Cdk1/CyclinB | pRS425-pGAL1 | DOM0071  (Gift from David Morgan) | Yeast | 3FLAG, M2 affinity | Gift from Skotheim Lab | Topacio et al., 2019 |
| Cdk2/CyclinA2/Cks2 | pETM41 | Civ Cells  (Gift from Barford Lab) | E.coli | Ni-NTA, TeV and 3C digest, S200 | Gift from David Barford  Co-expression | ^77^ |
| APPBP1/Uba3 | pGSTHs APPBP1 rbs UBA3 | BL21(DE3)  RIL | E.coli | Ni-NTA, Thrombin digest, MonoQ, S200 | Co-expression | Huang et al., 2005 |
| Ubc12 | pDEST17-UbE2M | BL21(DE3)  RIL | E.coli | Ni-NTA, MonoQ, S200 |  | Huang et al., 2005 |
| Cdc43b | pET11 His TEV Cdc34 | BL21(DE3)  RIL | E.coli | Ni-NTA, TeV Digest, MonoQ, S200 |  | ^63^ |
| Ubch5 | pGEX TEV UbcH5c | BL21(DE3)  RIL | E.coli | Ni-NTA, MonoQ, S200 |  | ^63^ |
| NEDD8 | pET11 His-Nedd8 | BL21(DE3)  RIL | E.coli | Ni-NTA, MonoQ, S75 |  | ^63^ |
| GSK3b | pET-GST | BL21(DE3)  RIL | E.coli | GST, TeV Digest, S200 |  | ^81^ |
| CK1a | pET-GST | BL21(DE3)  RIL | E.coli | GST, TeV Digest, MonoQ,  S200 |  | ^24^ |

Figure S2

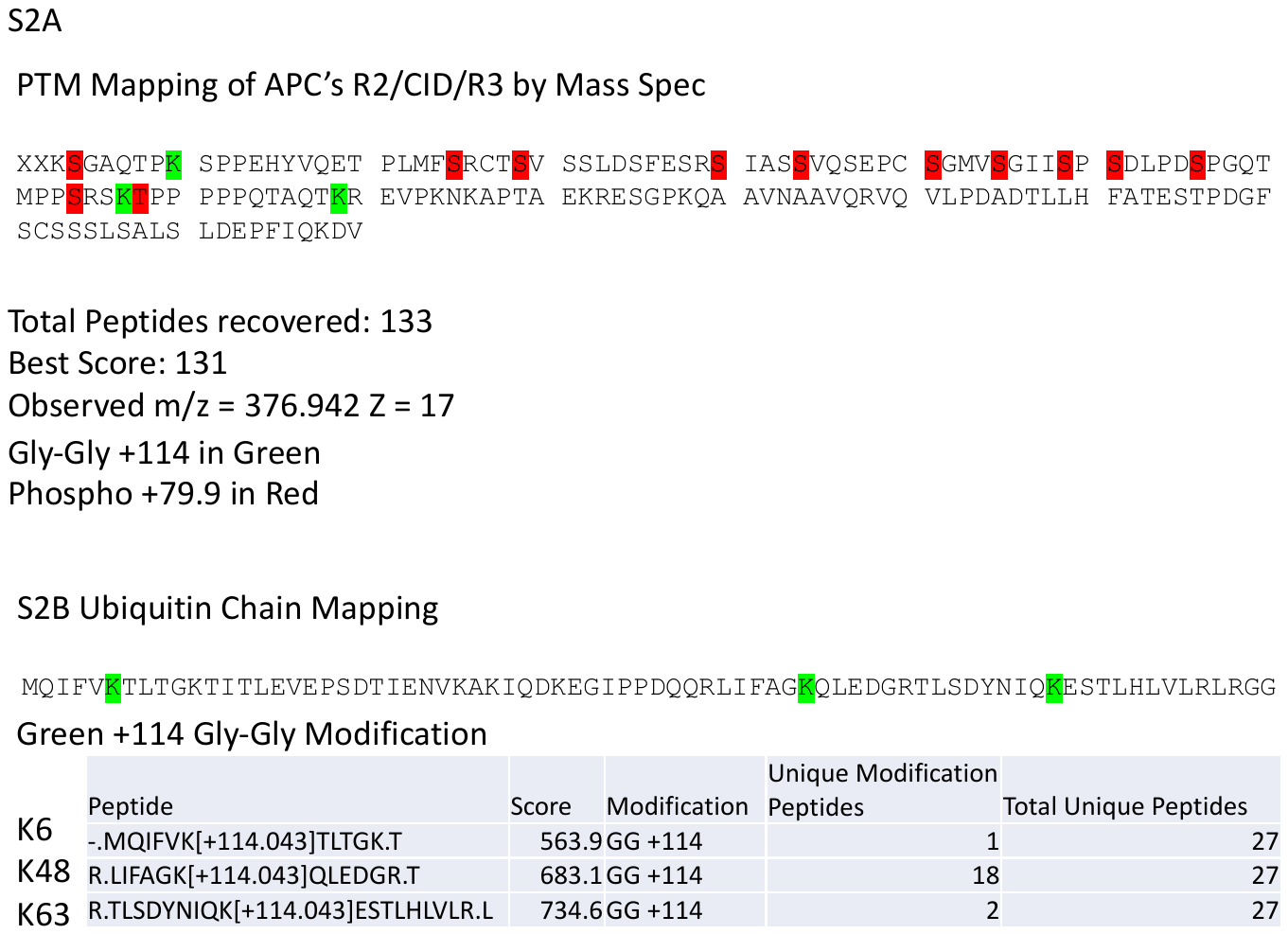

Figure 2SA: Mapping of post translational modificiations of APC’s R2/CID/R3 region by mass spec. Phospho-peptides identified by Byonic 2.14.2. Previous data from original data analysis in SEQUEST.

Figure 2SB: Mapping of ubiquitin chaining by LC-MS/MS PTM analysis for GG addition to lysines (+114Da).

Figure S3

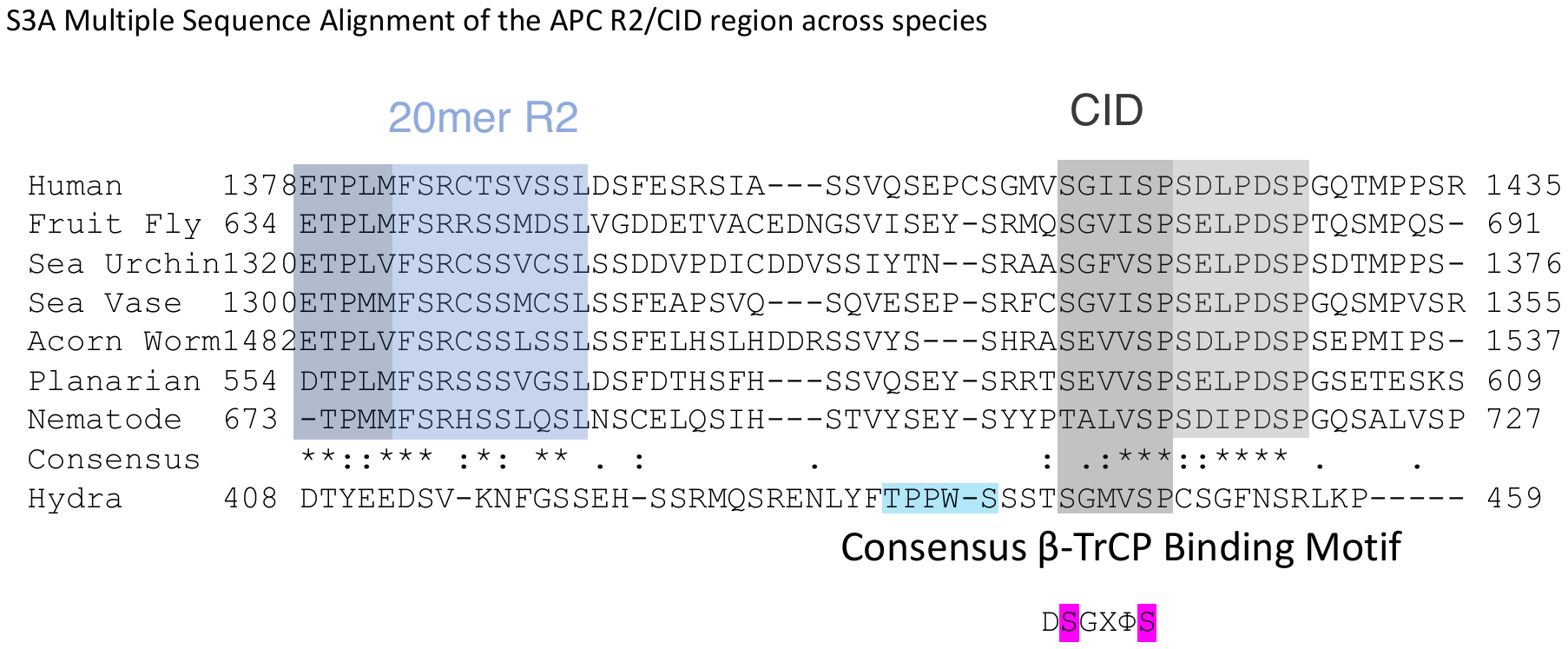

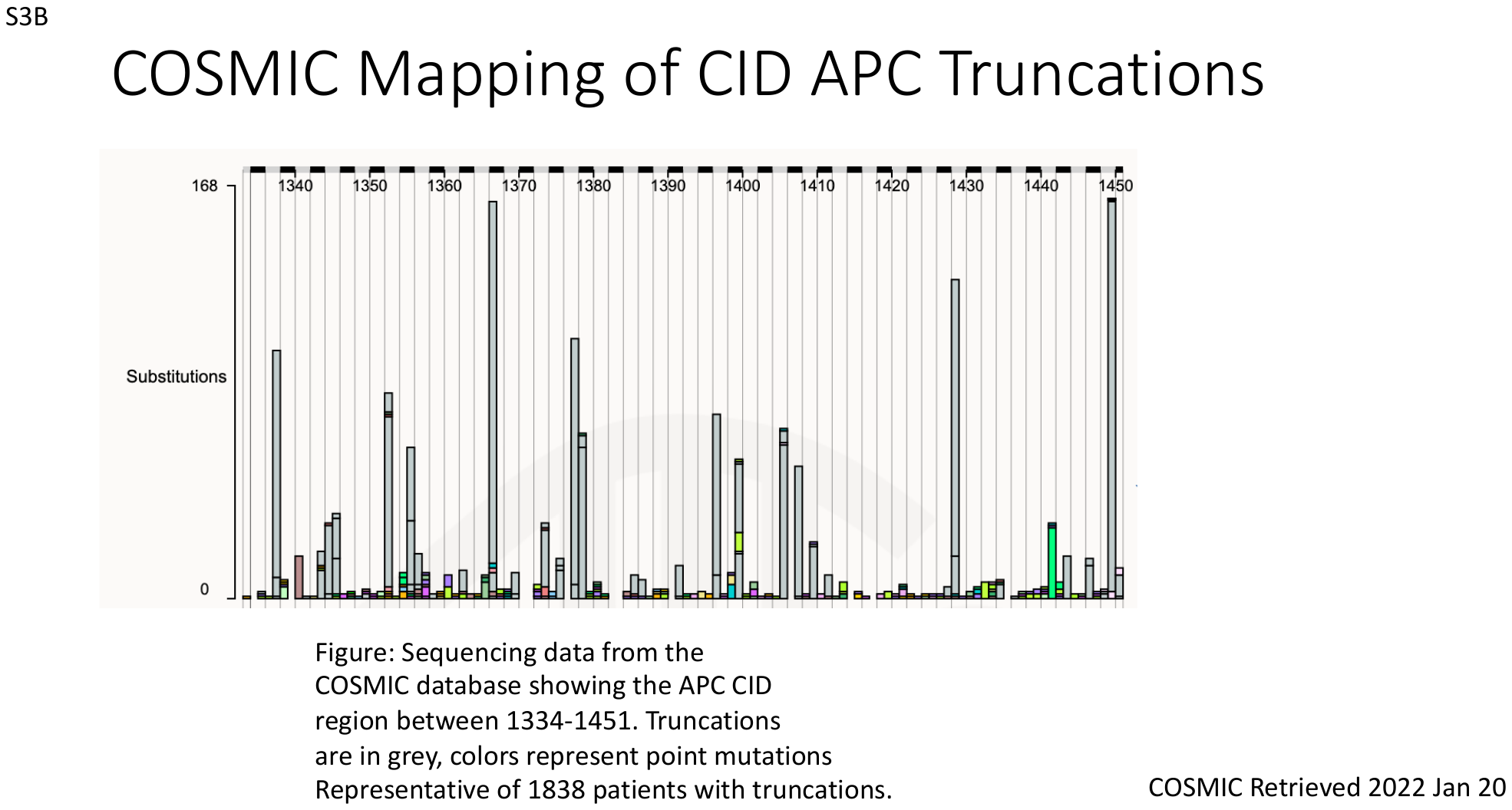

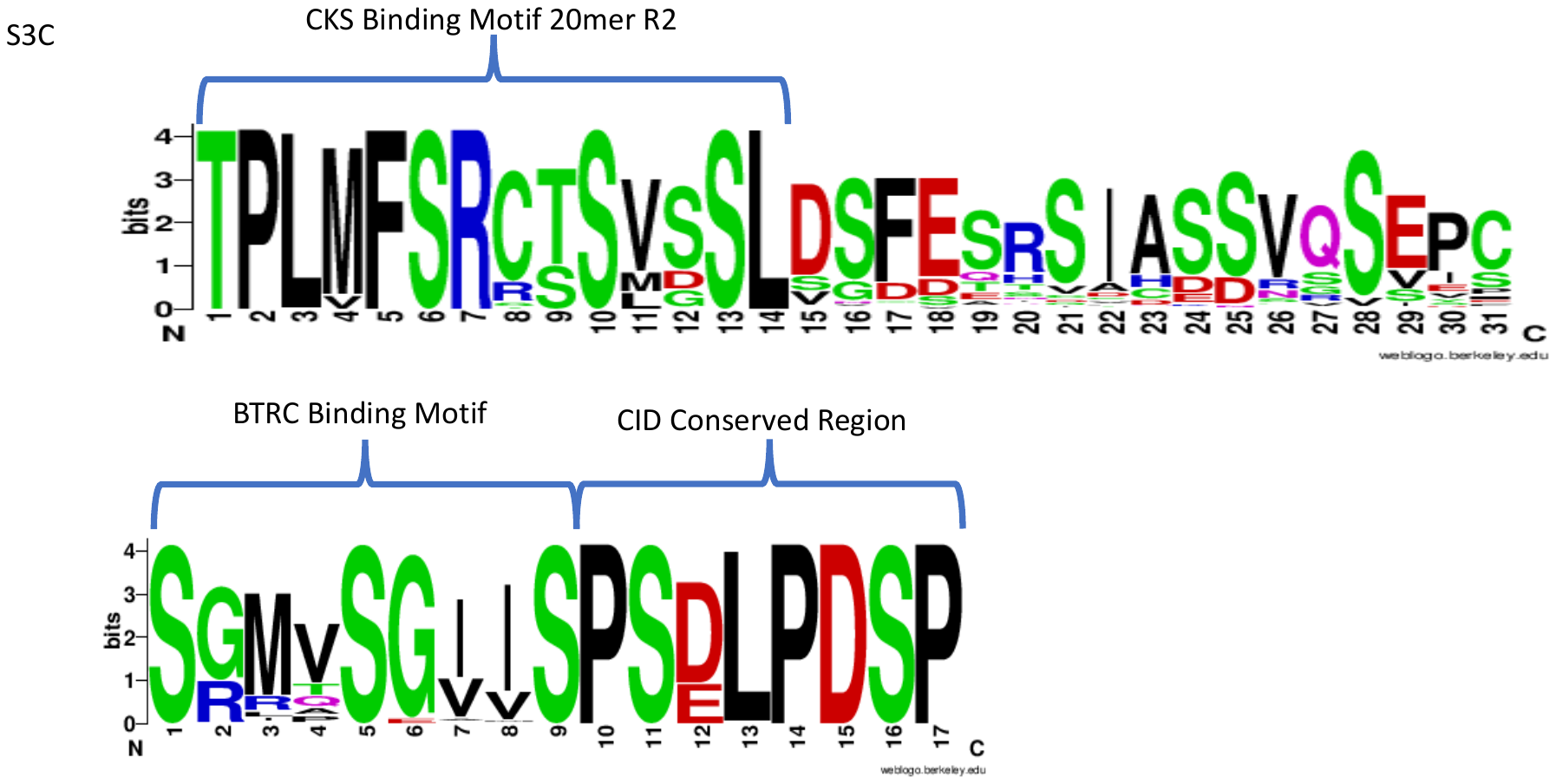

Figure S3A: Genetic analyses of APC R2/CID region on an evolutionary basis across species. This revealed that both the BTRC motif is highly conserved in most known organisms with Wnt.

Figure S3B: Mapping of APC CID region (1334-1451) truncations from the COSMIC database for 1838 patients.

Figure S3C: Motif conservation of 111 species of higher organisms showing extreme conservation of the Cks, 20mer R2 and BTRC binding motifs.

Figure S4

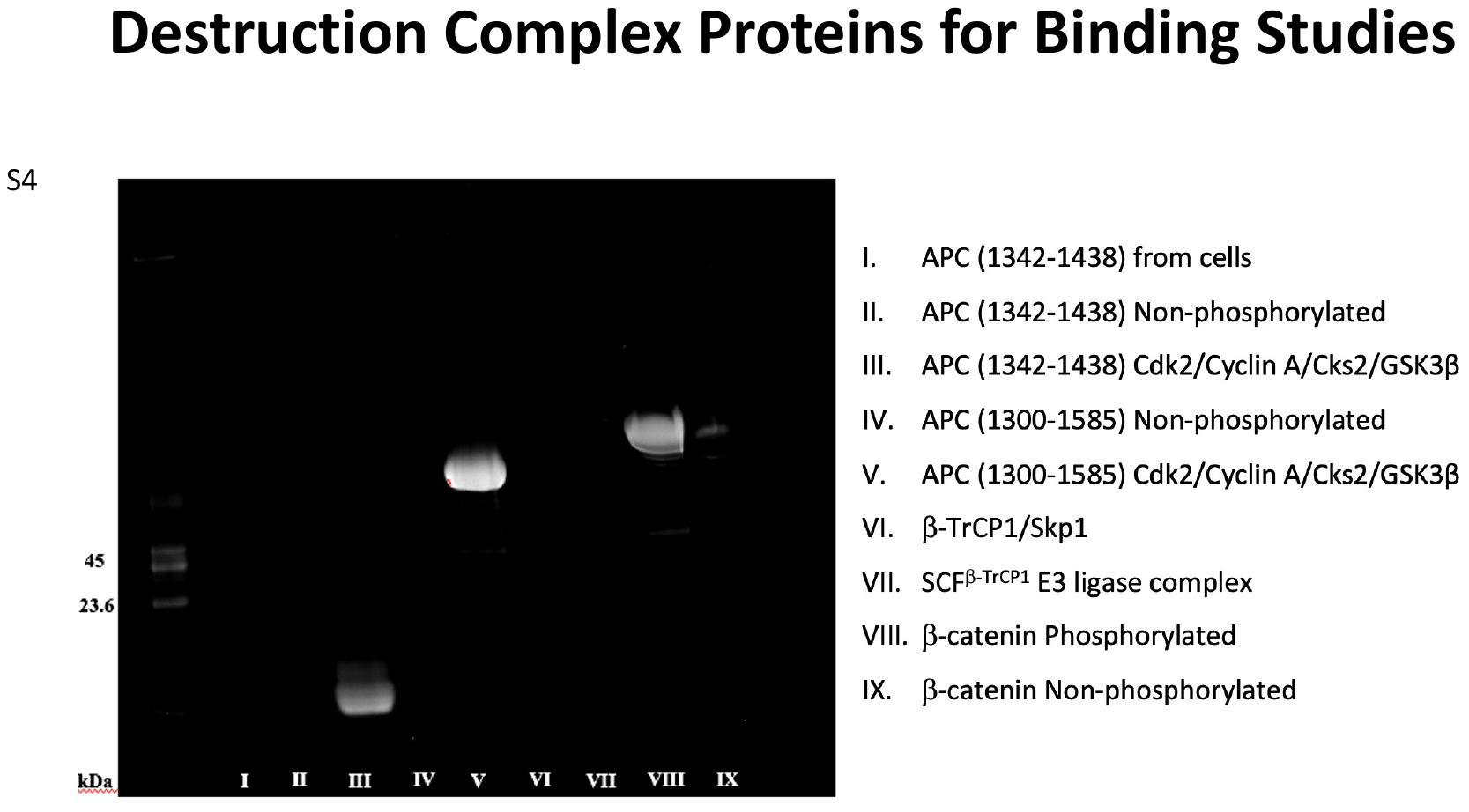

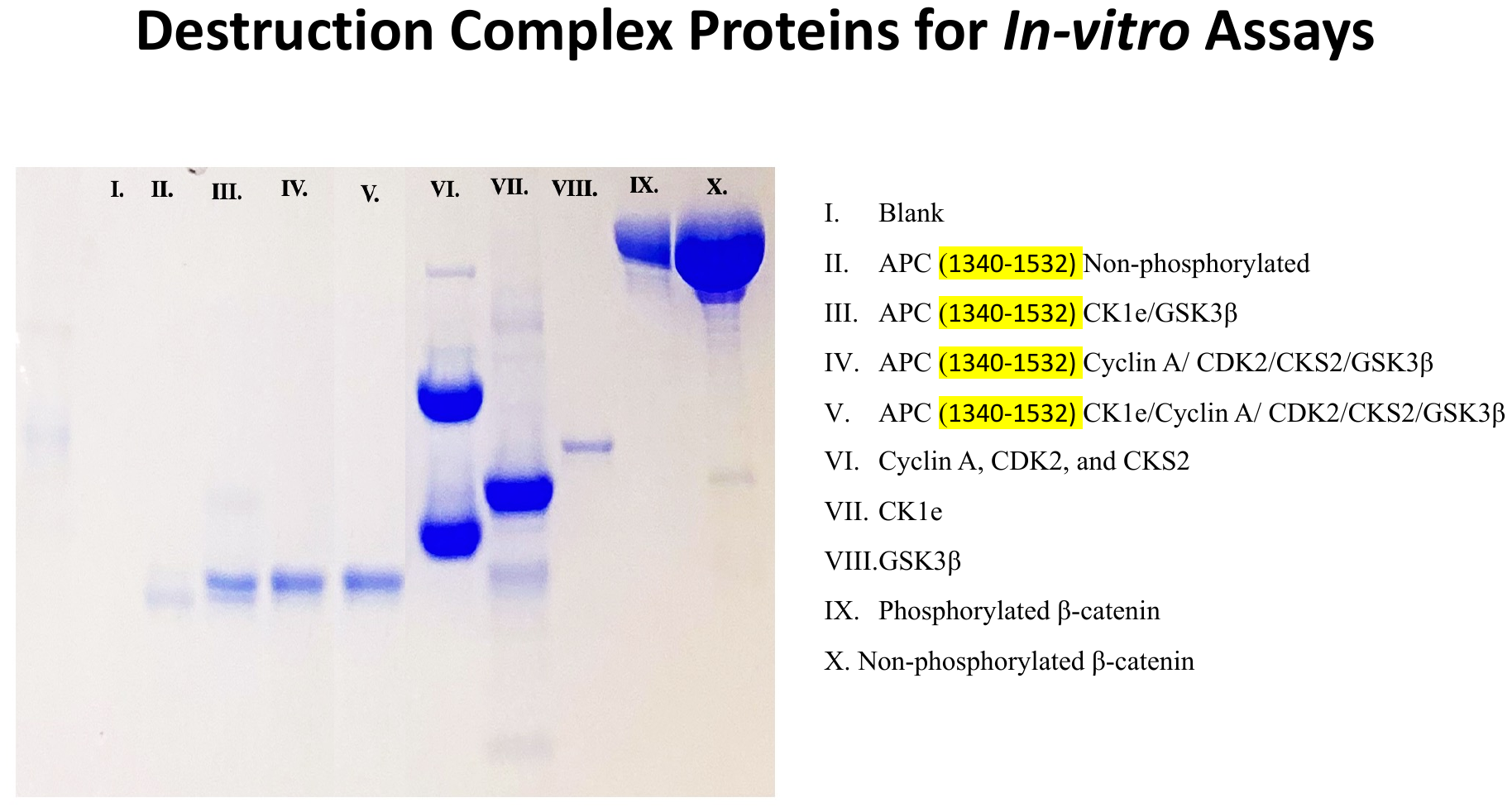

Figure S4: Phosphorylation of DC components analyzed on ProQ Phospho-stain gel (a) and Coomassie (b) for binding studies.
